# Cell cycle-coupled CK1δ turnover, autoinhibition, and activity

**DOI:** 10.64898/2026.03.18.712569

**Authors:** Fidel E. Serrano, Bianca Ruppert, Axel C. R. Diernfellner, Michael Brunner

## Abstract

Casein kinase 1δ (CK1δ) is a ubiquitously expressed kinase involved in diverse cellular processes, including cell cycle regulation. CK1δ activity is attenuated by (auto)phosphorylation. However, inhibitory phosphorylation is efficiently opposed by cellular phosphatases as CK1δ accumulates in its hypophosphorylated, active state. CK1δ is a target of the nuclear ubiquitin ligase APC/C-CDH1, yet the kinase is apparently stable. Thus, the physiological relevance of CK1δ (auto)phosphorylation, autoinhibition, and regulated turnover has remained unclear. Here we show that CK1δ activity and abundance are coordinated in a cell cycle-dependent manner. During G1, assembled CK1δ kinase is stable while free active kinase is degraded. In S phase, unassembled CK1δ seems to be no longer degraded, likely to support functions in DNA damage signaling. Upon mitotic entry, the downregulation of phosphatases promotes CK1δ (auto)phosphorylation and consequent autoinhibition, thereby preserving a pool of kinase to rapidly reestablish the post-mitotic steady state.

## Introduction

Casein kinase 1 (CK1) is an evolutionarily conserved serine/threonine kinase found throughout the eukaryotic lineage^1^, with homologs in yeast^2^, fungi^3^, plants^4^, and animals^5^. CK1 isoforms act as effectors in a variety of cellular signalling pathways^6,7^. All members of the CK1 family share a highly conserved, structured catalytic domain and a divergent, largely disordered C-terminal tail^8,9^. CK1 enzymes are phosphate-directed kinases displaying high affinity for motifs pre-phosphorylated (primed) at the-3 and to a lesser extent also the -4 position with the consensus pS/T-X-X-(X)-S/T but display much lower activity towards unprimed sites^10–12^. This specificity is conferred by a dedicated phosphate-binding pocket that correctly orients the primed substrate^11^. CK1s are considered constitutively active because they do not have regulatory subunits and are not activated by phosphorylation of their activation loop^13^.

Among the seven CK1 isoforms in humans (α, β, γ1, γ2, γ3, δ, and ε), CK1δ and CK1ε play crucial roles in regulating the circadian clock^5,10,12^, cytoskeletal organization^14,15^, cell cycle^16,17^, and development^18–20^. CK1δ and CK1ε activity is attenuated by phosphorylation of their C-terminal tail^8,9^. Autoinhibition occurs via a competitive mechanism in which the phosphorylated tail blocks the active site^21,22^. Inhibitory tail phosphorylation is mediated by autophosphorylation and phosphorylation by other cellular kinases^7,12^. Although C-terminal tail phosphorylation inhibits CK1δ/ε in vitro, the kinases undergoes futile cycles of phosphorylation and dephosphorylation^23^, keeping the kinase in a dephosphorylated, active state^24^. CK1δ/ε activity is controlled by several mechanisms, including sequestration^7^, leaving the physiological significance of this inhibitory mechanism unclear.

CK1δ is shuttling between nucleus and cytoplasm and localizes dynamically to the centrosome and pericentrosomal region^15,25,26^. Catalytically inactive CK1δ mutants accumulate in the nucleus and nuclear export of CK1δ depends on kinase activity^27,28^. CK1δ is a target of the nuclear ubiquitin ligase anaphase-promoting complex/cyclosome in association with its coactivator CDH1 (APC/C-CDH1)^29^, which is active from late anaphase/telophase through G1 and inactivated when the cell cycle progresses into S phase^30^. Yet, CK1δ appears to be stable^31,32^, raising the question about the physiological conditions under which APC/C-CDH1 targets the kinase.

We have recently analyzed the posttranslational regulation of CK1δ in unsynchronized cells, which are mainly in the G1 phase of the cell cycle^28^. These findings indicate that CK1δ is stabilized through assembly with binding partners such as the circadian clock protein PER2. In contrast, unassembled CK1δ, which accumulates at detectable levels upon overexpression, is rapidly degraded. This suggests that APC/C-CDH1 selectively targets the unassembled kinase, thereby limiting aberrant CK1δ activity. Here, we expand our analysis of CK1δ regulation in G1 and investigate how CK1δ stability and localization are controlled across the cell cycle.

## Results

### Subcellular distribution of CK1δ is dynamic

We have recently shown that CK1δ expressed at physiological levels is stable while overexpressed CK1δ which is not assembled with binding partners such as PERIOD2, is rapidly degraded, preferentially in the nucleus in a manner dependent on its kinase activity^28^. To characterize in more detail the subcellular dynamics, synthesis and turnover of endogenous CK1δ we performed immunofluorescence (IF) of U2OStx cells. In untreated cells, CK1δ was highly concentrated in the pericentrosomal region (Figs. 1A, C), consistent with its reported localization at the centrosome and Golgi apparatus^7,33^, and was also detectable in the nucleus, consistent with its reported association with PERIOD proteins^32^.

**Figure 1.**
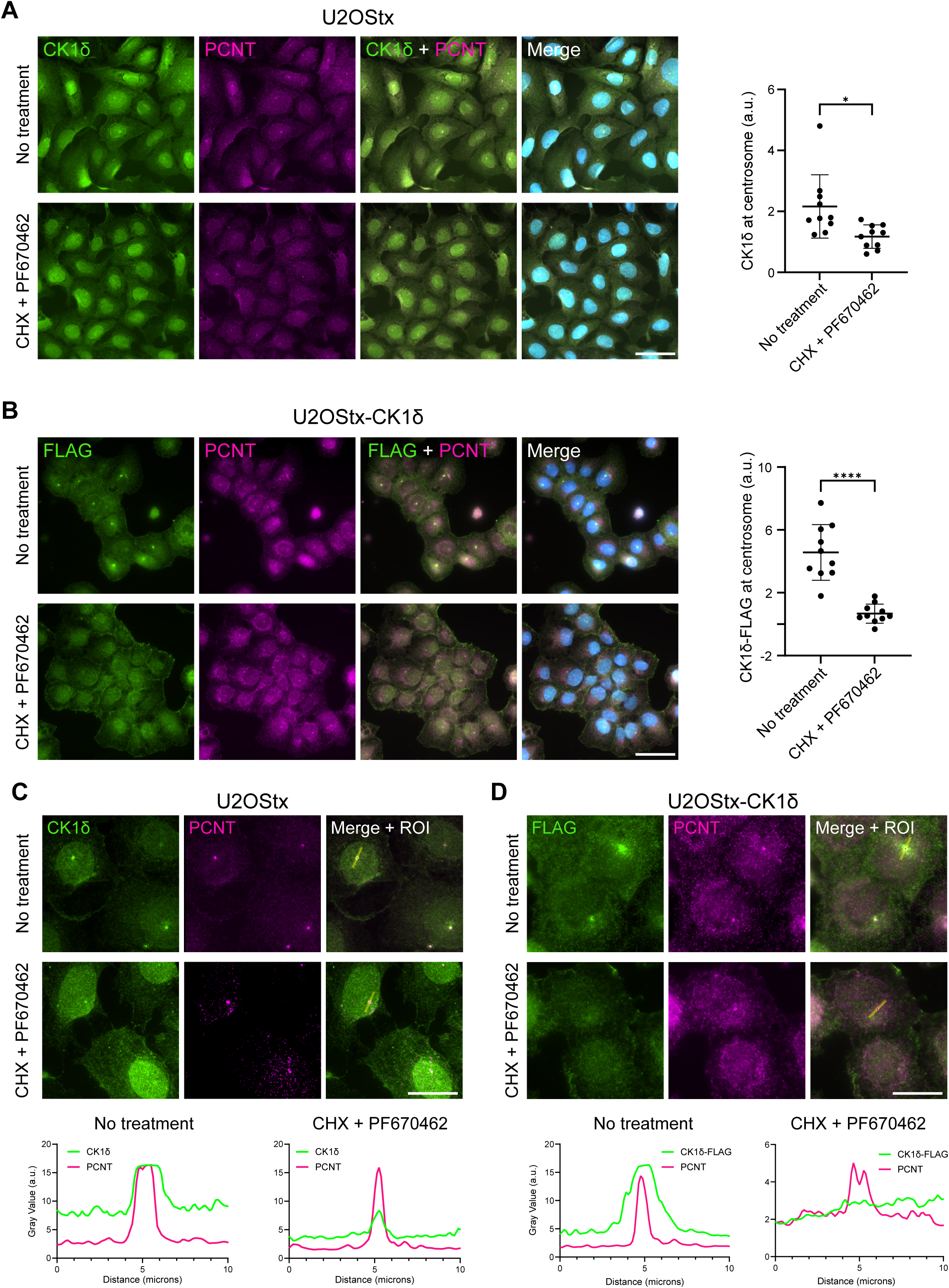
CK1δ inhibition results in decreased centrosomal localization. (A) U2OStx cells were treated with a combination of CHX and PF670462 for 4 h to inhibit protein synthesis and kinase activity or left untreated. Endogenous CK1δ and pericentrin (PCNT) were detected by IF. Green (AF488): anti-CK1δ, magenta (Cy3): anti-PCNT. Nuclei were identified with DAPI. In untreated cells, CK1δ exhibited clear nuclear and pericentrosomal staining. In CHX + PF670462-treated cells, pericentrosomal localization of CK1δ was decreased and CK1δ accumulated in the nucleus. Scale bar = 50 µm. Right panel: Density above surrounding background of CK1δ at pericentrosomal region. Quantification of n = 10 cells, Welch’s t-test. * indicates p-value < 0.05. (B) DOX-induced U2OStx_CK1δ cells were either left untreated or treated with CHX and PF670462 for 4 h and analyzed as described above. Green (AF488): CK1δ-FLAG was detected with anti-FLAG antibody. Magenta (CY3): anti-PCNT. In untreated cells, CK1δ-FLAG was concentrated at the pericentrosomal region. In CHX + PF670462-treated cells, pericentrosomal localization of CK1δ-FLAG was decreased and CK1δ-FLAG accumulated in the nucleus. Right panel: Density above surrounding background of CK1δ-FLAG at pericentrosomal region. Quantification of n = 10 cells. Welch’s t-test. **** indicates p-value < 0.0001. U2OStx cells (C) and DOX-induced U2OStx_CK1δ cells (D) were analyzed as described in (A) and (B), respectively. Fiji-based multi-channel line plot analysis showing the fluorescence intensity profiles of CK1δ (C) and CK1δ-FLAG (D) relative to PCNT along the indicated region-of-interest (ROI) in representative untreated cells and cells treated with CHX or PF670462. Scale bar = 20 µm.

We then asked whether the subcellular localization of CK1δ is static or dynamic. It has been shown that catalytically inactive CK1δ mutants accumulate in the nucleus^27^ and we have previously shown that kinase inhibition with PF670462, a specific CK1δ/ε inhibitor^34^, inhibits nuclear export of CK1δ and, in addition, protects the kinase from degradation. Therefore, to trap and stabilize shuttling CK1δ in the nucleus, U2OStx cells were treated for 4 h with PF670462 together with cycloheximide (CHX) to distinguish relocalization from de novo protein synthesis. Under these conditions, the localization of CK1δ at the pericentrosomal region was reduced and the kinase accumulated in the nucleus (Figs. 1A, C). These data demonstrate that the pericentrosomal localization of CK1δ is dynamic and in equilibrium with the nuclear pool. A fraction of CK1δ remained associated with the pericentrosomal region after the 4 h treatment with PF670462 (Fig.1 C).

Next, we analyzed U2OStx cells stably overexpressing doxycycline (DOX)-inducible FLAG-tagged CK1δ. Expression of CK1δ-FLAG was induced for 24 h with DOX to generate a pool of unassembled kinase and analysed by IF. Despite overexpression, the subcellular distribution of CK1δ-FLAG was similar to that of CK1δ in control cells, displaying pronounced concentration of CK1δ-FLAG in the pericentrosomal region (Fig. 1B). Treatment with PF670462 resulted in nuclear accumulation of CK1δ-FLAG and reduction of pericentrosomal staining (Figs. 1B, D). These data indicate that pericentrosomal and nuclear CK1δ exist in a dynamic equilibrium, regardless of whether the kinase is expressed at physiological levels or overexpressed.

Thus, at steady state, the subcellular levels of endogenous or overexpressed CK1δ are not static but are dynamically maintained by the balance between subcellular shuttling, binding equilibria with the pericentrosomal region. We have recently shown that unassembled CK1δ is rapidly degraded in the nucleus^28^. Our data suggest that under physiological expression levels, rebinding of CK1δ to the pericentrosomal region constitutes the dominant pathway, effectively outcompeting nuclear degradation, while overexpression promotes degradation of free nuclear CK1δ. Such a mechanism would enable continuous monitoring and regulation of CK1δ levels while maintaining a low abundance of unassembled, potentially harmful kinase. Because free CK1δ levels are kept low and pericentrosomal binding sites are not saturated^28^, the pericentrosomal CK1δ pool may function as a buffered reservoir that can supply newly emerging binding partners such as PERIOD proteins, when they accumulate at higher levels than free CK1δ.

To test this hypothesis, we employed U2OStx cells stably expressing mKate2-tagged Cryptochrome 1 (mK2-CRY1). These U2OStx-mK2-CRY1 cells were transfected with PER2 to rapidly overexpress a binding partner of CK1δ. As recently shown^28^, overexpressed PER2 and mK2-CRY1 mutually stabilize each other and co-accumulate in nuclear foci, whereas in untransfected cells mK2-CRY1 alone is unstable, expressed at a low level and homogeneously distributed throughout the nucleus (Fig. 2A). Thus, the formation of nuclear mK2-CRY1 foci serves as an indicator of PER2 transfection^28^. Because U2OStx cells exhibit low transfection efficiency, only a fraction of U2OS-mK2-CRY1 cells expressed PER2. This allowed a side-by-side comparison between endogenous CK1δ in PER2-expressing cells identified by nuclear mK2-CRY1 foci from non-expressing cells within the same population. In untransfected cells, endogenous CK1δ was concentrated at the single centrosome (Fig. 2B, red arrows). In contrast, in PER2-transfected cells, CK1δ accumulated at elevated levels and colocalized with mK2-CRY1 in PER2-dependent nuclear foci (Fig. 2B). Notably, all nuclear CK1δ foci were mK2-CRY1 positive, whereas CK1δ foci lacking mK2-CRY1, which would indicate pericentrosomal localization of CK1δ, were not detected in PER2-transfected cells. This change in CK1δ localization upon formation of nuclear mK2-CRY1 foci is consistent and can be seen in multiple fields-of-view (FOV). Together, these findings suggest that PER2 overexpression competes for CK1δ binding and, when expressed at very high levels, can even displace CK1δ from the pericentrosomal region.

**Figure 2.**
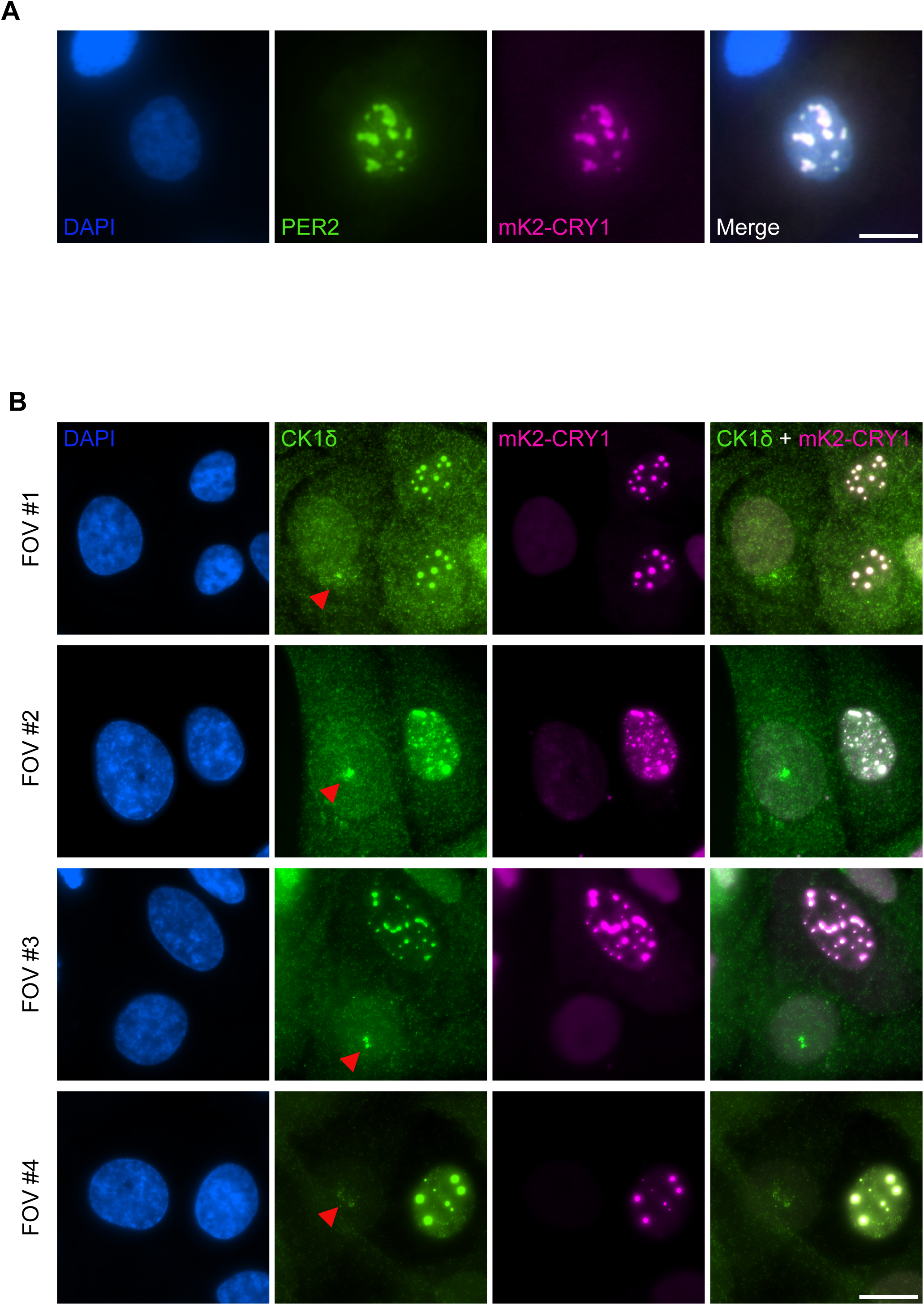
PER2 overexpression displaces CK1δ from the centrosome. U2OStx_mK2-CRY1 cells were transfected with PER2, followed by DOX induction. (A) IF for PER2 shows the formation of nuclear foci containing both overexpressed PER2 and mK2-CRY1 as reported previously^28^. Scale bar = 20 µm. (B) IF for CK1δ shows that the kinase expressed from its endogenous locus is recruited to the nuclear PER2-CRY1 foci and depleted from the pericentrosomal region, in agreement with previous observations^28^. Four FOVs are shown. Presence and absence of nuclear mK2-CRY1 foci indicate PER2-transfected and untransfected cells, respectively. Pericentrosomal CK1δ localization, accessed by CK1δ positive (green) and CRY1 negative (magenta) foci, was observed in all untransfected cells examined (50 out of 50 cells evaluated). In PER2 and CRY1-expressing cells CK1δ accumulated in CRY1-positive (magenta) nuclear foci. No cell (0 out of 15 cells evaluated) displayed pericentrosomal localization of CK1δ (CRY1-negative focus). Fisher’s exact test, p = 4.8 × 10⁻¹□. Scale bar = 20 µm.

### Dephosphorylation of CK1**δ** by cellular phosphatases

Phosphorylation of CK1δ at the kinase C-terminal tail can be stabilized and detected by treating cells with phosphatase inhibitors^35^. We compared the kinetics of tail phosphorylation upon treatment of cells with Calyculin□A (CalA), a potent inhibitor of the protein phosphatases PP1 and PP2A, and okadaic acid (OA), which inhibits PP2A more effectively than PP1^36^. For this purpose, we used U2OStx stably expressing under a doxycycline (DOX)-inducible promoter either FLAG-tagged wild-type CK1δ or its catalytically inactive (kinase-dead) version, CK1δ-K38R^12^. In U2OStx_CK1δ cells, both CalA and OA inhibited CK1δ dephosphorylation and led to the accumulation of elevated levels of hyperphosphorylated CK1δ (Fig. 3A), suggesting that phosphorylation protects the overexpressed kinase from degradation. CalA does not inhibit PP2C or protein tyrosine phosphatases and is largely ineffective against PP2B and PP7. Thus, the sensitivity of CK1δ dephosphorylation to CalA suggests PP1, PP2A, PP4, and PP5 as likely candidates contributing to dephosphorylation^36^. Although sensitivity to CalA and OA cannot rigorously distinguish among these phosphatases, the high OA concentrations required to inhibit CK1δ dephosphorylation suggest the involvement of PP1, which is 10 - 100-fold less sensitive to OA than PP2A, PP4, or PP5^35,36^. Thus, CK1δ is actively maintained in a dephosphorylated state, most likely through the contribution of PP1, with possible additional contributions from PP2A, PP4, and PP5. PP1 has previously suggested as a major phosphatase of CK1δ/ε^37^. Notably, PP1, PP2A, and PP4 activities are downregulated during mitosis^38,39^, pointing to a potential physiological role for CK1δ tail phosphorylation and autoinhibition during this stage of the cell cycle.

**Figure 3.**
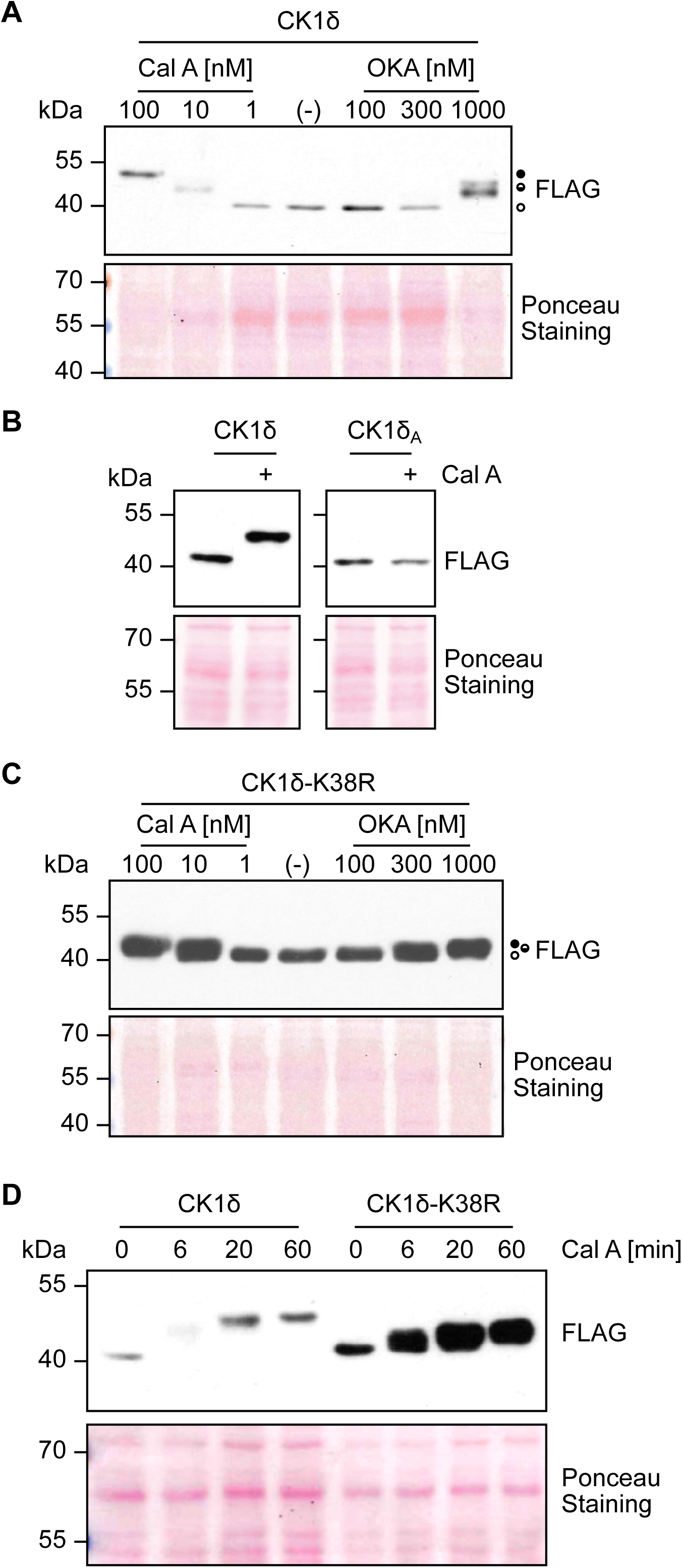
Phosphatase inhibition results in CK1δ phosphorylation. (A) Immunoblot analysis of CalA-and OA-induced phosphorylation of CK1δ-FLAG. Phosphorylation of CK1δ is strongly induced by CalA and, albeit with markedly lower efficiency, also by OA. n = 3. (B) Immunoblot analysis demonstrating the absence of a CalA-induced electrophoretic mobility shift in the CK1δ_A_ mutant kinase. n = 3. (C) Immunoblot analysis of CalA-and OA-induced phosphorylation of kinase-dead CK1δ-K38R. n = 3. (D) Time-course of CalA-induced phosphorylation of CK1δ and CK1δ-K38R. n = 3. Ponceau S staining of membranes are shown as loading control.

To analyze whether the electrophoretic mobility shift induced by CalA treatment was indeed caused by phosphorylation of the CK1δ tail, we generated a U2OStx cell line expressing a CK1δ variant in which all serine and threonine residues in the C-terminal tail were replaced by alanine (CK1δ_A_). Treatment of these cells with CalA did not induce an electrophoretic mobility shift of the tail mutant kinase (Fig.□3B).

To separate autophosphorylation from phosphorylation by other kinases, we examined U2OStx_CK1δ-K38R cells. Upon CalA or OA treatment, the kinase-dead CK1δ-K38R accumulated at elevated levels and was also phosphorylated but to a lesser extent than wild-type CK1δ (Fig.□3C), suggesting that fewer phosphorylation sites are targeted by other kinases. Because CK1δ-K38R is already rather stable^40^, the data suggest that partial tail phosphorylation is sufficient to further stabilize CK1δ-K38R. Time-course experiments revealed that CK1δ became hyperphosphorylated within 1□h of CalA treatment, with intermediate phosphorylation states visible at 6 and 20□min (Fig.□3D), in agreement with previous observations^8^. CalA-induced phosphorylation of CK1δ-K38R followed similar kinetics (Fig.□3D). We noted that treatment with a PPase inhibitor led to increased levels of overexpressed, and thus unassembled, CK1δ (Fig. 3A, D) as well as CK1δ-K38R (Fig. 3C, D), suggesting that full or partial phosphorylation of the CK1δ tail stabilizes both active and inactive forms of the kinase.

### Tail phosphorylation protects CK1**δ**/**ε** from degradation

To examine in more detail how tail phosphorylation affects the turnover of unassembled CK1δ, U2OStx cells stably expressing CK1δ-FLAG were either left untreated or pre-treated for 1 h with the phosphatase inhibitor CalA, alone or together with PF670462, followed by a 90-min cycloheximide (CHX) chase. In CHX-treated cells, overexpressed CK1δ was rapidly degraded, with a half-life of approximately 15 min (Fig. 4A), consistent with our previous findings^40^. The 1 h pre-treatment with CalA induced hyperphosphorylation of CK1δ, whereas co-treatment with PF670462 and CalA produced an intermediate phosphorylation state (Fig. 4A), similar to the CalA-induced phosphorylation of CK1δ-K38R, indicating that PF670462-insensitive kinases targeted a fraction of the phosphorylation sites in CK1δ tail. Both 1 h pre-treatments resulted in elevated CK1δ expression levels, indicating that most of the newly synthesized overexpressed kinase was degraded in untreated cells. This conclusion was confirmed by the CHX chase. Hyperphosphorylated CK1δ remained stable throughout the CHX chase (Fig. 4A, C), indicating that tail phosphorylation protects the overexpressed kinase from rapid degradation. CK1δ shares a high degree of sequence identity and potential functional redundancy with CK1ε, with both kinases implicated in circadian clock regulation^5,41^□and Wnt signalling^42^. Treatment of cells overexpressing FLAG-tagged CK1ε with CHX, CalA, and PF670462 revealed that CK1ε turnover is also regulated in a similar manner to CK1δ (Fig.□4B,□D).

**Figure 4.**
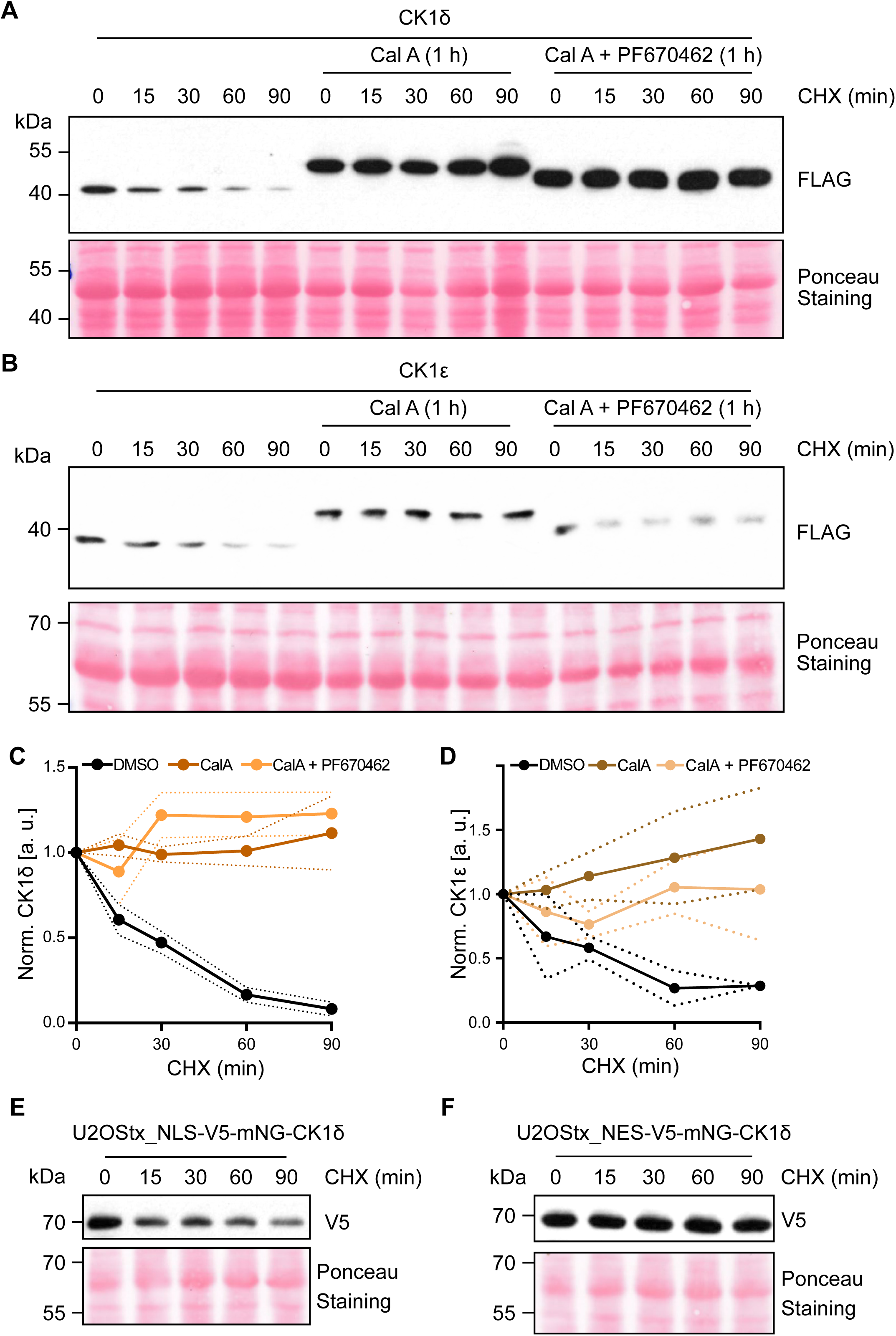
CK1δ/ε phosphorylation stabilizes overexpressed kinase. (A) Phosphorylation protects overexpressed CK1δ from degradation. U2OStx_CK1δ cells were incubated with cycloheximide (CHX) (left part) or pretreated for 1 h with CalA (middle part) or with CalA + PF670462 (right part). CK1δ was analyzed at the indicated time points after CHX addition. Overexpressed CK1δ is rapidly degraded. CalA and CalA + PF670462 induce maximal and intermediate phosphorylated CK1δ and stabilize the kinase. n = 3. (B) CalA and CalA + PF670462 induce phosphorylation of CK1ε and stabilize the kinase. n = 3. (C) Densitometric quantification of n=3 Western blots (see A) shown as mean ± SD. Overexpressed unphosphorylated CK1δ is degraded with a half-life of about 15 min. Both CalA and CalA + PF670462 treatments stabilize the kinase. (D) Densitometric quantification of n=3 Western blots (see B) shown as mean ± SD. (E) CK1δ is preferentially degraded in the nucleus. U2OStx cells stably expressing NLS-mNG-CK1δ and NES-mNG-CK1δ were incubated with CHX. Samples were analyzed at the indicated time points after CHX addition. Overexpressed NLS-mNG-CK1δ is unstable while NES-mNG-CK1δ is stable. Ponceau S staining of membranes are shown as loading control.

CK1δ is a target of the nuclear ubiquitin ligase APC/C-CDH1^29^. To directly analyze the subcellular turnover of CK1δ, we employed U2OStx cell lines expressing mNeonGreen-tagged CK1δ fused either to an SV40 NLS or to an HIV-1 NES, which direct the kinase predominantly to the nucleus or cytosol, respectively^40^. Western blot analysis of cycloheximide-treated cell lines revealed that NLS-mNG-CK1δ was degraded more rapidly than NES-mNG-CK1δ.

### Cell cycle-dependence of CK1**δ** expression and phosphorylation

Because CK1δ has various functions at different stages of the cell cycle, including S phase and mitosis^6,17^, we investigated whether expression and phosphorylation state of CK1δ are regulated in a cell cycle-dependent manner. CK1δ is a target of the nuclear ubiquitin ligase APC/C-CDH1^29^, which is active from late anaphase/telophase all the way through G1 and inactive in S phase, G2 and mitosis^30^. To analyze whether inactivation of APC/C-CDH1 affects expression levels of unassembled CK1δ, U2OStx_CK1δ were arrested at G1/S with a double thymidine block. The block was then relieved and CK1δ expression was analyzed (Fig. 5A). CK1δ levels increased after release from the G1/S arrest, compared to non-arrested cells (which are predominantly in G1) and to cells arrested at the G1/S transition (Fig. 5B). The accumulated CK1δ remained dephosphorylated, indicating that the phosphatases acting on the kinase tail were active during S phase. These findings suggest that overexpressed, and thus unassembled CK1δ accumulates in a dephosphorylated, active state and escapes nuclear degradation at cell cycle states when APC/C-CDH1 is inactive. The data suggest that inactivation of APC/C-CDH1 in S-phase may protect free nuclear CK1δ to facilitate its functions in DNA damage repair and checkpoint signalling.

**Figure 5.**
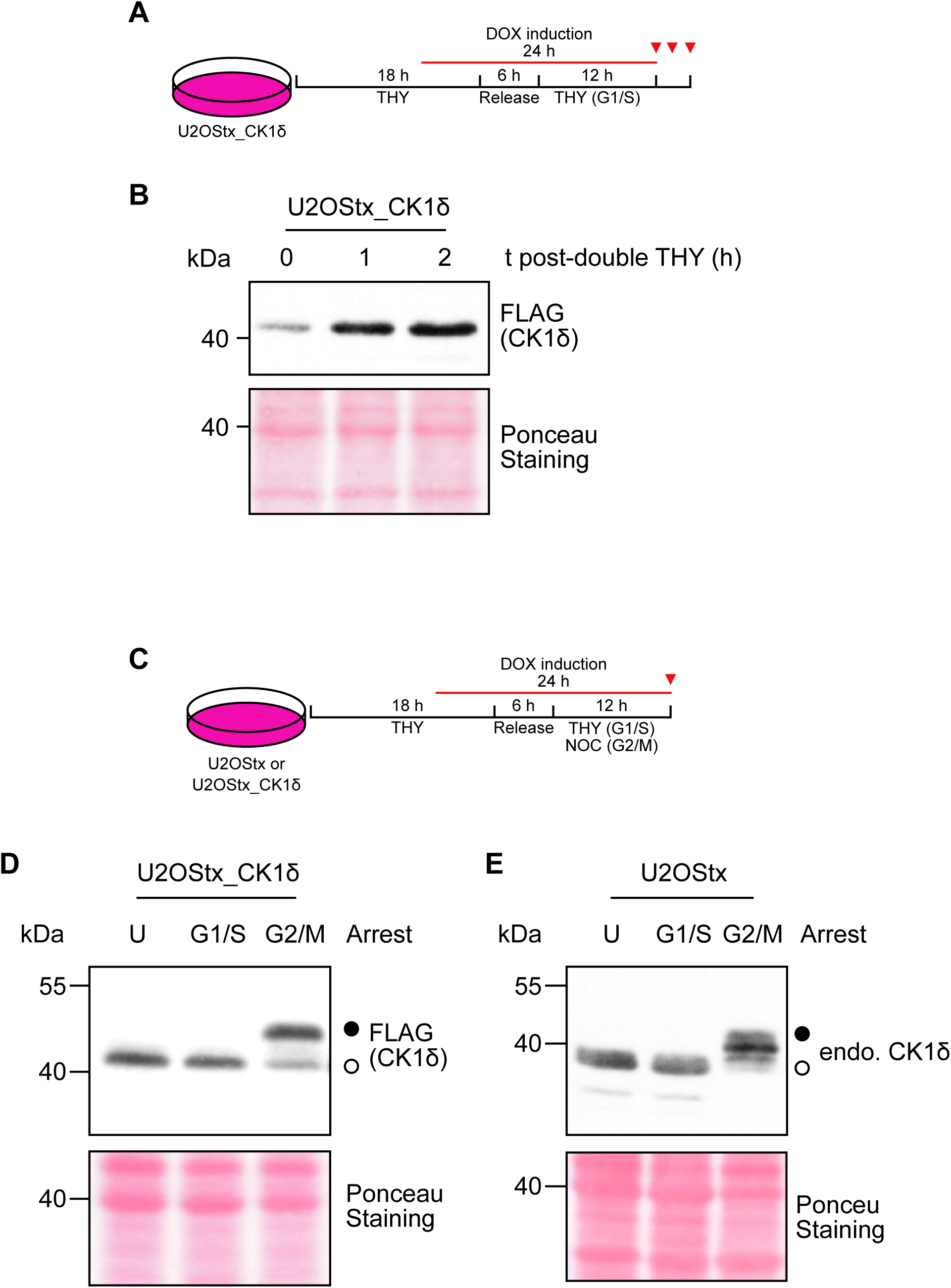
CK1δ is stabilized post-S phase and phosphorylated during G2/M. (A) Schematic of S phase arrest and release of U2OStx_CK1δ cells. (B) Overexpressed CK1δ accumulates post-S phase release. n = 3. (C)) Schematic of G1/S and G2/M phase arrest of U2OStx_CK1δ or U2OStx cells (D, E) Overexpressed CK1δ in U2OStx_CK1δ cells (D) and endogenous CK1δ in U2OStx cells (E) are unphosphorylated in unsynchronized and G1/S-arrested cells. G2/M-arrested cells accumulate hyperphosphorylated CK1δ. n = 3. Ponceau S staining of membranes are shown as loading control.

Upon entry into mitosis, the activity of major cellular phosphatases such as PP1 and PP2A is globally attenuated^38,39^, while distinct pools of these phosphatases remain active at defined subcellular structures^43^. We reasoned that this inhibition might facilitate the accumulation of phosphorylated CK1δ. To test this, we arrested U2OStx_CK1δ cells at the G1/S boundary using a double thymidine block, and at G2/M using a thymidine-nocodazole block followed by mitotic shake-off to enrich for cells in mitosis^44,45^ (Fig. 5C). CK1δ-FLAG was then analyzed via immunoblotting. In unsynchronized (U) and G1/S-arrested cells, CK1δ was hypo-or unphosphorylated. In contrast, G2/M-arrested cells accumulated tail-phosphorylated CK1δ (Fig. 5D). Similarly, endogenous CK1δ was hyperphosphorylated upon G2/M arrest of U2OStx control cells while it remained hypophosphorylated in unsynchronized and G1/S-arrested cells (Fig. 5E).

To assess the localization of CK1δ during mitosis, we performed IF on U2OStx_CK1δ cells staining for the overexpressed kinase. Based on the DAPI staining we assigned the fixed cells to distinct cell-cycle stages. In G2/prophase, identified by duplicated centrosomes and increased DNA content, CK1δ localized predominantly to structures which correspond to the duplicated centrosomes (Fig. 6A, 1^st^ column), consistent with previous reports^15,25^. In metaphase and anaphase (Fig. 6A, 2^nd^ column), characterized by DNA condensation, overexpressed CK1δ localized to centrosomes and polar microtubules, consistent with previous reports on the localization of endogenous kinase^14^. In addition, a considerable fraction of the overexpressed CK1δ was evenly distributed throughout the cell, suggesting that unassembled CK1δ was not targeted for degradation. During telophase and cytokinesis, as chromosome segregation and cell division progressed, CK1δ was diffusely distributed across the dividing cell (Fig. 6A, 3^rd^ column). Although it is not possible to identify cells that have only recently exited mitosis and entered G1 in fixed samples, all G1 cells displayed a concentration of CK1δ at the pericentrosomal region (Fig. 6A, 4^th^ column). Similar results were observed when using U2OStx cells and staining for endogenous CK1δ (Fig. 6B).

**Figure 6.**
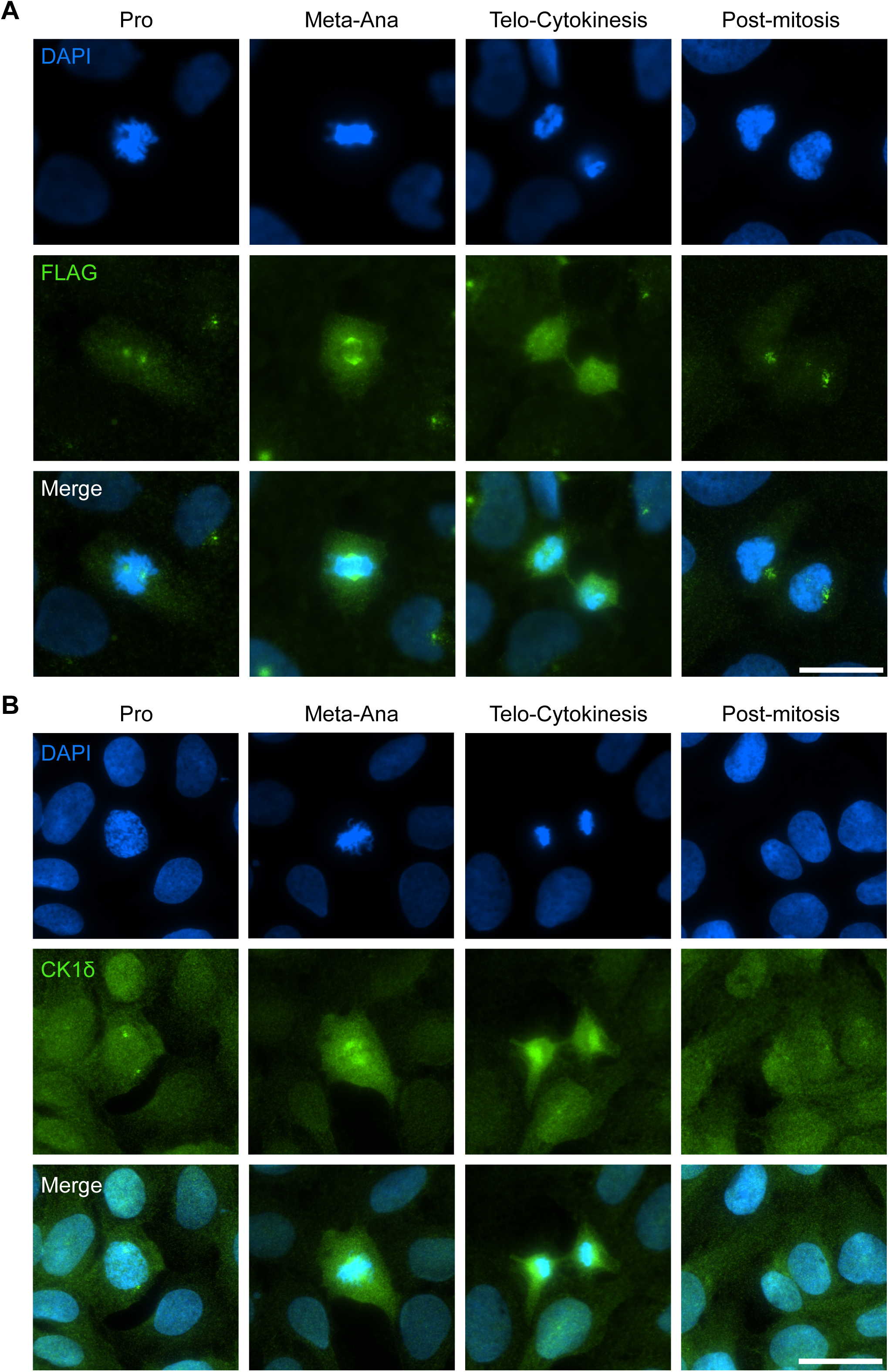
CK1δ localization during mitosis. (A) Upper panels: DAPI-staining of U2OStx_CK1δ cells was used to identify stages of the cell cycle. Middle panels. IF analysis of overexpressed CK1δ with FLAG antibody. Lower panels: Merged DAPI and FLAG image. Scale bar = 20 μm. (B) Upper panels: DAPI-staining of U2OStx cells was used to identify stages of the cell cycle. Middle panels. IF analysis of endogenous CK1δ with anti-CK1δ antibody. Lower panels: Merged DAPI and CK1δ image. Scale bar = 20 μm. Pro: Prophase, Meta-Ana: Metaphase-Anaphase, Telo-Cytokinesis: Telophase-Cytokinesis.

## Discussion

A central and long-standing question in the regulation of casein kinase 1δ (CK1δ) concerns the physiological role of its autoinhibitory C-terminal tail phosphorylation. Biochemical studies have established that autophosphorylation of the CK1δ tail suppresses catalytic activity, suggesting an intrinsic mechanism for kinase regulation. However, this model appears difficult to reconcile with observations in cells, where CK1δ is found predominantly in a dephosphorylated, and therefore catalytically active state. This discrepancy raises a general conundrum: if CK1δ is constitutively active in vivo, how is its activity effectively regulated, and under what physiological circumstances does tail-mediated autoinhibition become relevant?

A related paradox has emerged in the circadian clock field. CK1δ plays a central role in regulating the stability and nuclear function of circadian PER proteins and is therefore expected to be readily available to ensure robust circadian timekeeping. In principle, CK1δ should not be limiting for PER phosphorylation. Yet, overexpression of CK1δ consistently accelerates the circadian clock, implying that kinase availability can influence clock speed. This finding suggests that CK1δ activity may be regulated not only by catalytic mechanisms but also by its spatial availability within the cell.

We have recently begun to address this paradox by demonstrating the importance of subcellular localization as it relates to CK1δ function and activity. We have shown that CK1δ is not limiting in the cytosol but becomes limiting in the nucleus in the context of the circadian clock^28^. This present study provides a potential explanation for how cells reconcile a low synthesis rate of CK1δ and low levels of free kinase with the need for efficient association with binding partners such as PER proteins. Our data indicate that CK1δ is dynamically stored at the pericentrosomal region, potentially forming a buffering pool that can be readily mobilized to bind to its partners. Such a mechanism would allow the cell to maintain low levels of free, potentially harmful, active kinase while simultaneously ensuring that CK1δ is readily available to newly synthesized PER proteins. In this way, pericentrosomal sequestration could function as a buffering system that couples limited CK1δ production to the rhythmically high demand for PER binding and phosphorylation in circadian timekeeping.

Here, we have also addressed the fate of CK1δ across the cell cycle. Because the pool of endogenous free CK1δ is small due to slow synthesis and binding to the pericentrosomal region^28^, we employed overexpression to analyze localization and degradation of unassembled CK1δ. Our data indicate that unassembled CK1δ is degraded in G1, coinciding with cell cycle stages in which its nuclear ubiquitin ligase, APC/C-CDH1, is active^46,47^. We show that hyperphosphorylated CK1δ accumulated in G2/M-arrested cells under both endogenous expression and overexpression conditions. Integrating our findings with previous studies, we propose that in G1 (Fig. 7), CK1δ is maintained in an active dephosphorylated state. This active CK1δ is dynamically stored at the pericentrosomal region and unassembled molecules are eliminated by the proteasome, primarily in the nucleus through APC/C-CDH1^28,29^. Because the synthesis rate of CK1δ is low and nuclear kinase is readily exported to the cytosol where it binds to the centrosome and Golgi, the fraction of CK1δ targeted for nuclear degradation is small and not detectable against higher steady state levels. However, using CK1δ overexpression allows to further uncover this degradation pathway. In S phase, APC/C-CDH1 is inactivated by CDH1 phosphorylation and inhibition of the ubiquitin ligase by its pseudo-substrate^30,48^. Because phosphatases remain active during S phase, CK1δ is maintained in a dephosphorylated, active state. We propose that unassembled kinase is no longer degraded in the nucleus, likely allowing free kinase to support DNA repair and checkpoint control^14,17,49^. CK1 phosphorylation triggers β-TrCP-mediated degradation of MDM2 and activates p53, thereby enhancing p53-dependent responses involved in checkpoint signaling and DNA repair^50,51^. Following DNA repair, CK1δ promotes checkpoint exit by phosphorylating WEE1, leading to its β-TrCP-dependent degradation and consequent relief of CDK1 inhibition. CK1δ also supports CHK1 stability during replication stress and, together with CK1ε, phosphorylates Topo IIα to facilitate DNA decatenation in late S/G2, thereby contributing to the orderly resumption of cell-cycle progression^17,52^. Upon mitotic entry, downregulation of phosphatase activity including PP1 and PP2A^38,39^, and activation of mitotic kinases^53^ facilitate (auto)phosphorylation and autoinhibition of CK1δ, terminating its DNA repair functions. Because CK1δ also contributes to mitotic processes^14,17,26,54,55^, a fraction of CK1δ assembled with its binding partners may remain active through locally retained phosphatase activity^43^. The majority of CK1δ, however, becomes phosphorylated and stabilized in an autoinhibited form, safeguarding it for transiently from degradation at mitotic exit, when both phosphatases and APC/C-CDH1 regain activity. At the onset of G1, reactivated CK1δ redistributes preferentially to the pericentrosomal region, while excess dephosphorylated kinase is degraded. By reactivating the safeguarded pool of CK1δ, cells can rapidly re-establish G1-specific steady-state levels without entirely replenishing the kinase through de novo synthesis, which is rather slow. We have no data regarding the order of events or their kinetics after cells exit mitosis. In particular, we do not know how rapidly CK1δ is dephosphorylated by reactivated phosphatases nor how efficiently it associates with pericentrosomal binding sites to escape degradation by reactivated APC/C-CDH1. Furthermore, our data do not address whether and how tail phosphorylation regulates CK1δ in G1. However, the bulk of CK1δ remains dephosphorylated in G1 indicating that tail phosphorylation is transient. Together, our findings reveal that the activity and abundance of dephosphorylated and phosphorylated CK1δ are regulated in a cell cycle-dependent manner (Fig. 7), suggesting distinct physiological functions associated with each phosphorylation state of the kinase. Although our model integrates the new findings reported here together with the data on the regulation of CK1δ in G1^28^ and the broader body of knowledge regarding both CK1δ biology and cell-cycle regulation, it is far from complete, and many important questions remain open. However, the model offers plausible novel concepts and mechanistic ideas that provide a basis for further investigation and discussion.

**Figure 7.**
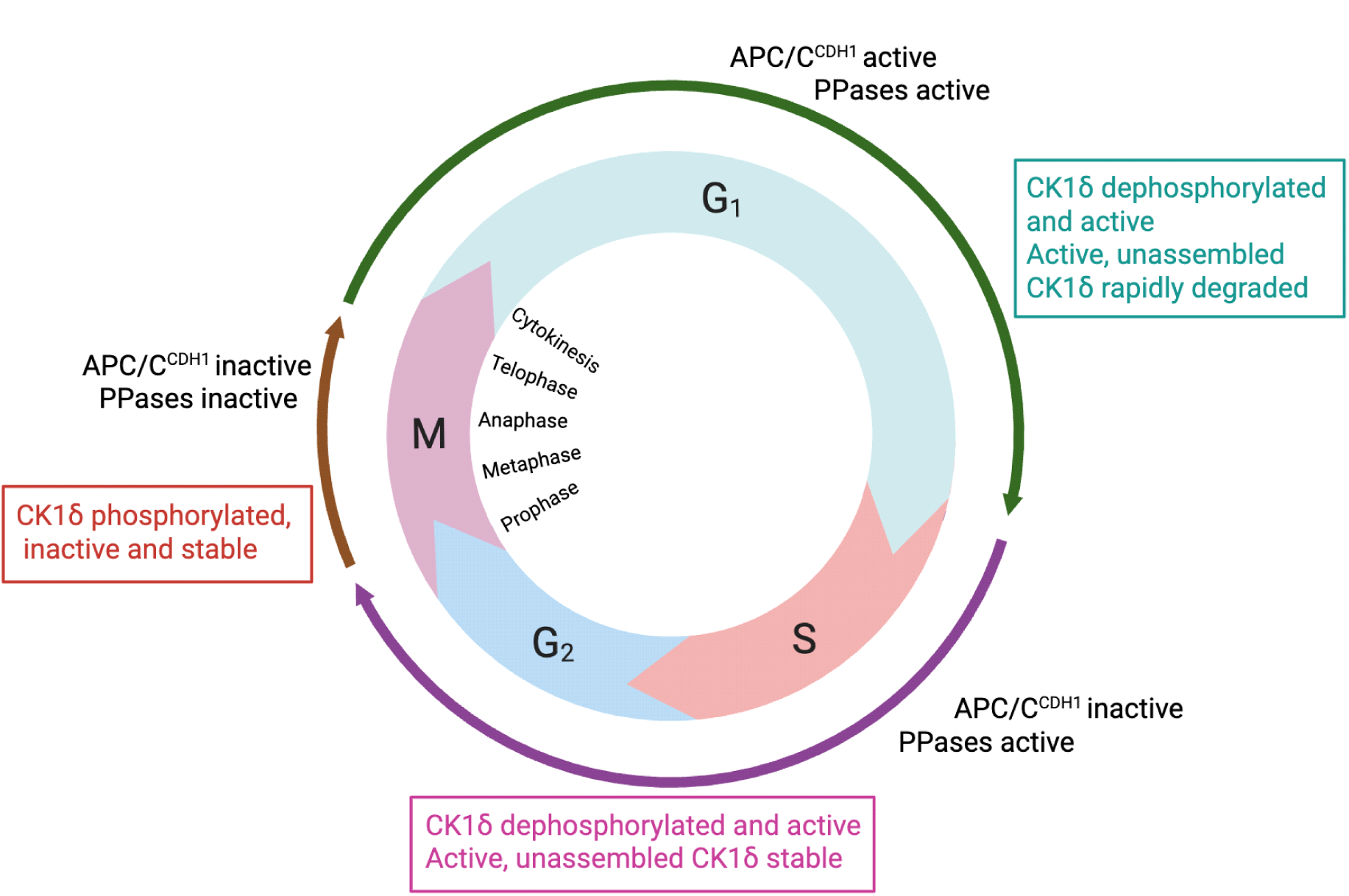
Model of CK1δ regulation over the course of the cell cycle. For details see main text (discussion section).

## Materials and Methods

### Key resources table

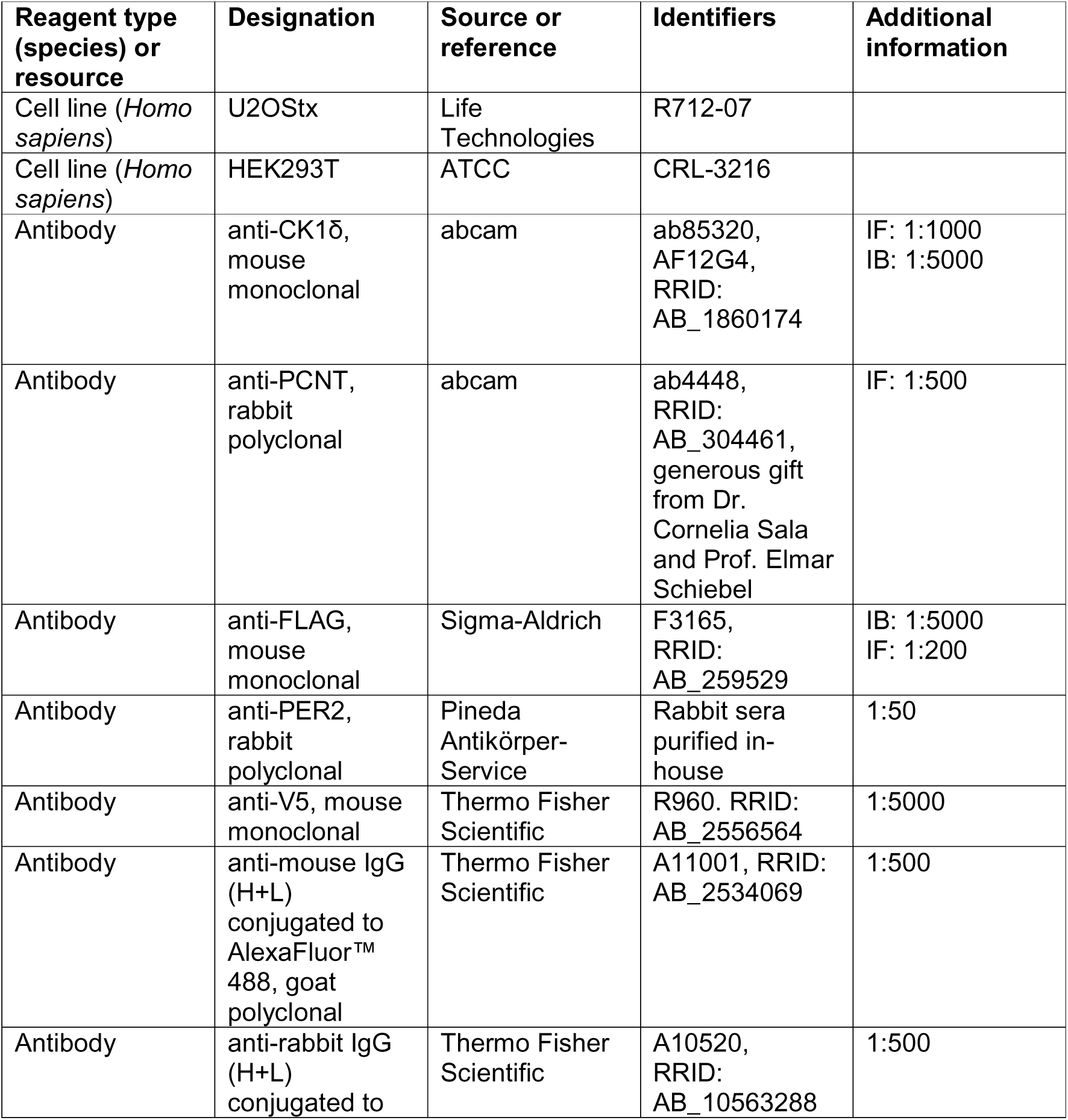
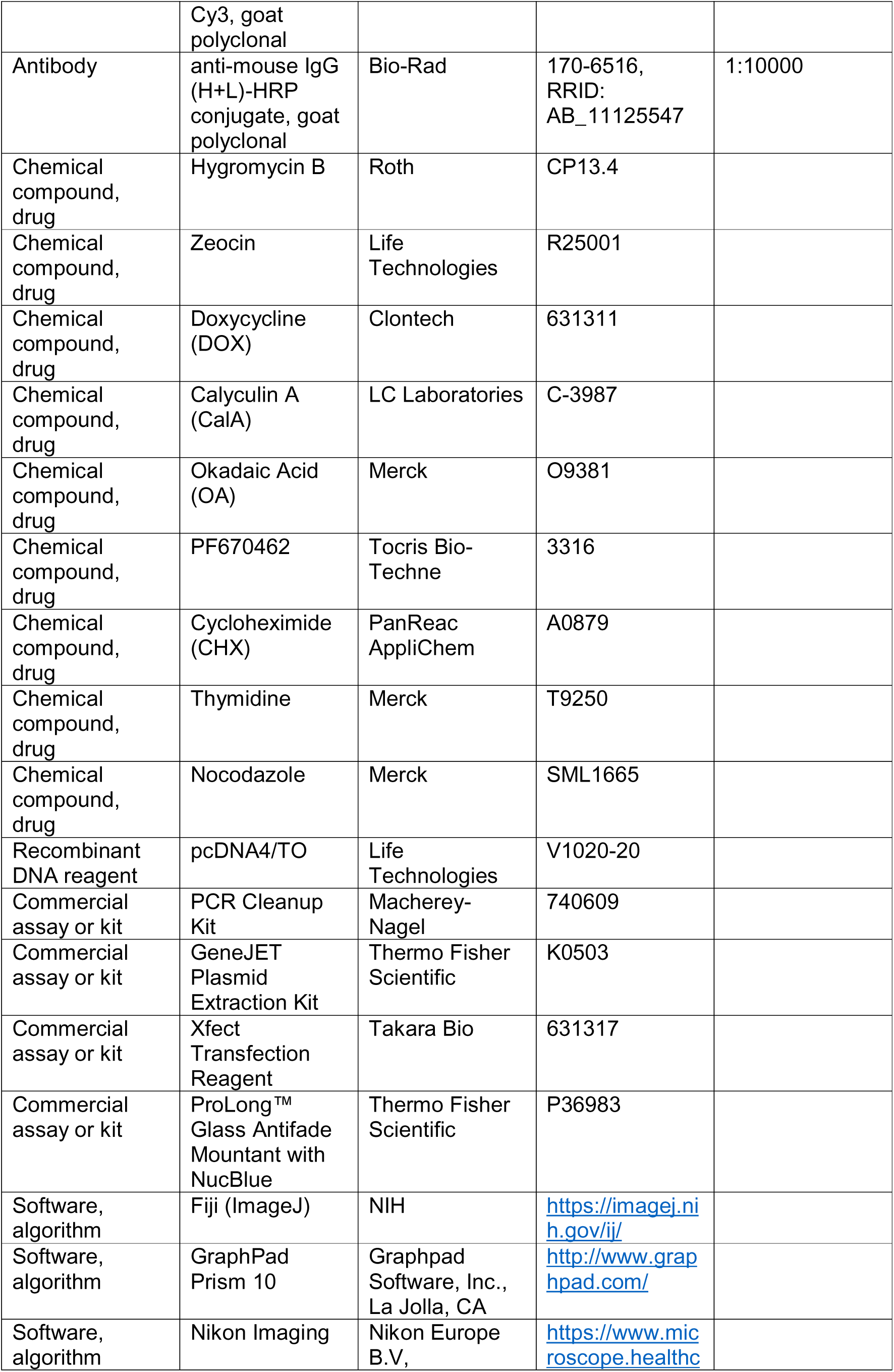
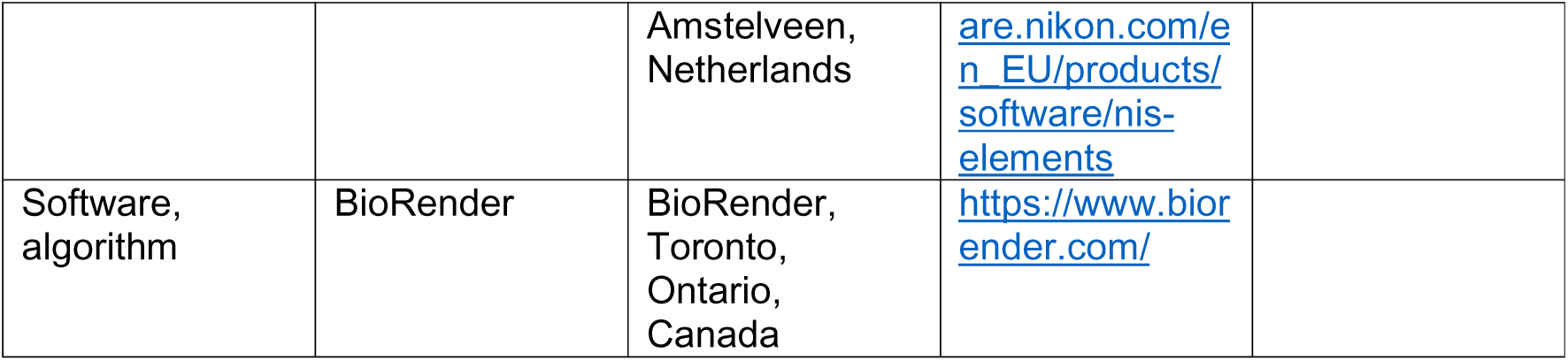

### Plasmids

Plasmids were constructed as previously described^12^. Briefly, restriction digestion, dephosphorylation of vector backbone and ligation was performed as recommended by New England Biolabs (NEB). Linear DNA was purified using NucleoSpin Gel and PCR Clean-up kit from Macherey-Nagel. Plasmids were extracted with GeneJET Plasmid Miniprep kit from Thermo Fisher Scientific. For cloning with overlapping regions (restriction enzyme-free cloning), enzymes (Q5 DNA polymerase, restriction enzyme DpnI and T4 DNA polymerase) were obtained from NEB. DpnI digestion and exonuclease reaction with T4 DNA polymerase was performed in NEBuffer 2.1. *E. coli* were transformed with exonuclease treated PCR product(s) to facilitate *in vivo* plasmid assembly via overlapping DNA ends. pcDNA4/TO-derived vectors were constructed by cloning the coding regions of the human CK1δ1 splice variant, referred to as CK1δ throughout the manuscript, and CK1ε, downstream of the CMV-TetO2 promoter which allows for DOX-inducible protein expression in cells harboring the tetracycline-repressor cassette (tx; i.e., U2OStx) and constitutively high expression in cells without the tetracycline-repressor cassette (i.e., HEK293T). Mutations (CK1δ1-K38R) and C-terminal FLAG-tags were introduced via ligation-free overlap cloning. All plasmid DNA sequences were sequence-verified prior to use.

### Cell culture and stable cell line generation

T-REx-U2OS (U2OStx; Life Technologies) and HEK293T (ATCC) cells were maintained in Dulbecco’s Modified Eagle Medium (DMEM) supplemented with 10% Fetal Bovine Serum (FBS) and 1x Penicillin-Streptomycin (Pen-strep). Cell culture reagents were obtained from Life Technologies. Cells were grown and maintained at 37°C in a humidified incubator containing 5% CO2.

For plasmid transfection, U2OStx cells were transfected with Xfect Transfection Reagent (Takara Bio Inc., Cat# 6313317) following the manufacturer’s protocol whereas HEK293T cells were transfected with polyethylenimine (PEI, made in-house) following the conventional protocol. To generate stable U2OStx cell lines, basal U2OStx cells were seeded in a 24-well plate and grown to confluence overnight. Cells were transfected with pcDNA4/TO vectors containing coding regions of CK1δ, CK1δ-K38R, or CK1ε with C-terminal FLAG epitopes. Stable transfectants were selected by growing cells to sub-confluence in complete media supplemented with 50 µg/mL hygromycin and 100 µg/mL zeocin (Invitrogen, Cat# R25001) over the course of two to three weeks. Resulting single cell colonies after selection were isolated for further experimentation as previously described. To induce protein overexpression, cells were treated with 10 ng/mL DOX.

### Immunofluorescence

U2OStx, U2OStx_CK1δ, or U2OStx_mK2-CRY1 cells were seeded on 12mm #1.0 glass coverslips (VWR, CAT# 631-1577P). Immunofluorescence was done either post-seeding or post-plasmid transfection and 24 h post DOX-induction. Cells were washed with PBS and fixed with 4% paraformaldehyde for 10 min at room temperature. Cells were permeabilized with PBS + 0.1% Triton X-100 for 10 min, then washed three times and blocked with PBS + 5% heat-inactivated FBS for 1 h.

Cells were then treated with either mouse monoclonal anti-CK1δ (abcam ab85320, clone AF12G4, 1:5000) or mouse monoclonal anti-FLAG antibody (Sigma-Aldrich F3165, 1:200), or rabbit polyclonal anti-PCNT (abcam ab4448, generous gift from Dr. Cornelia Sala and Prof. Elmar Schiebel, 1:500), or rabbit polyclonal anti-hPER2 (in-house, 1:50) diluted in PBS + 3% BSA + 0.25% Tween20 for 1 h to detect endogenous or overexpressed CK1δ, respectively. This was followed by washing with PBS, and treatment with the corresponding secondary antibody goat anti-mouse conjugated to AlexaFluor488 (AF488; Thermo Fisher Scientific A11001, 1:500) or goat anti-rabbit conjugated to Cy3 (Thermo Fisher Scientific A10520, 1:500) diluted in PBS + 3% BSA + 0.25% Tween20 for 1 h at 37°C. Cells were again washed three times, then mounted onto glass slides with ProLong™ Glass Antifade Mountant with NucBlue (Thermo Fisher Scientific P36983) and sealed with nail polish.

### Fluorescence microscopy

For microscopy of fluorescently-tagged proteins and IF, a widefield Nikon Ni-E microscope was used (Nikon Imaging Center, Heidelberg), at a 60X magnification oil immersion objective. The corresponding filter sets for DAPI, AF488, Cy3, and mK2 were then used to visualize the specific signal from each fluorophore used. Exposure times were optimized and kept constant within each experiment to obtain comparable fluorescence intensities between treatment conditions (i.e., no treatment vs. CHX + PF670462 treatment). For image acquisition, the central focal plane was determined and images were taken as at 0.3 µm z-steps to a total 11 steps.

### Digital image analysis

To analyze the resulting micrographs, built-in programs in FIJI were used. To quantify CK1δ localization at the centrosome, the PCNT channel was used to detect the subcellular localization of the centrosome. The signal intensity at the corresponding CK1δ and FLAG channels were then quantified and background-subtracted. These values were then tabulated for statistical analysis via Welch’s t-test using the using the built-in analysis in GraphPad Prism 10. The resulting data was visualized using GraphPad Prism 10.

To perform the line plot analysis, a region-of-interest (ROI) was selected using the line selection tool followed by using the built-in plot profile function in FIJI. These data were then visualized using GraphPad Prism 10.

### Immunoblotting

Protein extraction was performed using an in-house lysis buffer (25□mM Tris-HCl, pH 8.0, 150□mM NaCl, 0.5% Triton X100, 2□mM EDTA, 1□mM NaF, freshly added protease inhibitors: 1 mM phenylmethylsulfonyl fluoride (PMSF), leupeptin (5 μg/ml), and pepstatin A (5 μg/ml)) according to the conventional protocol. Briefly, cells were scraped and sample was collected via centrifugation. The sample was incubated in ice-cold lysis buffer on ice for 10 min and then incubated in a sonicated water bath for 10 min. Samples were centrifuged at 16,000 x g, 4 °C, 15 min taking the supernatant. Protein concentration was quantified via Nanodrop (NP80, Implen). For immunoblot analysis, protein samples were analyzed by SDS-polyacrylamide gel electrophoresis and transferred to a nitrocellulose membrane (cytiva, Amersham Protran, 0.45 µm pore size) via semi-dry blotting. To control for protein loading, membranes were stained with Ponceau S. Membranes were typically incubated in primary antibody at 4°C overnight and in secondary antibody conjugated to horseradish peroxidase at room temperature for 2 h. The following primary antibodies were used: anti-FLAG (Sigma-Aldrich F3165, 1:5000, mouse monoclonal), anti-CK1δ (abcam AF12G4, 1:5000, mouse monoclonal), and the following secondary antibody was used for luminescent detection: goat anti-mouse IgG (H+L)-HRP conjugate (Bio-Rad 170-6516, 1:10000). For decoration, membranes were incubated in decoration buffer (100 mM Tris-HCl pH 8.5, 2.2 × 10-2 % (w/v) luminol, 3.3 × 10-3 % (w/v) coumaric acid, 9.0 × 10-3 % (v/v) H2O2) and exposed for a time series to X-ray films (Fujifilm). Films were then developed (Medical film processor from Konica Minolta). Densitometric analysis of immunoblots was done using the built-in module in FIJI. The results were measured and plotted onto GraphPad Prism for downstream analysis (i.e., plotting densitometric signal over time post-CHX chase).

### Phosphatase inhibition and cycloheximide chase

To characterize phosphatase activity, cells were treated with either CalA or OA. To inhibit phosphatases over a time course, cells were treated with 80 nM CalA (LC Labs Cat# C-3987) for 60, 20, 6, and 0 (untreated) min. All chemical treatments were performed with the corresponding solvent (vehicle) control such as DMSO for CalA and H_2_O for CHX and PF670462.

Unless otherwise stated, induction of protein expression was typically performed by addition of 10 ng/mL doxycycline (DOX; Thermo Fisher Scientific Cat# 631311) 24 h prior to downstream experiments. To arrest protein translation for a cycloheximide (CHX) chase, CHX was added to our cells to a final concentration of 10 µg/mL. Untreated samples were taken as the 0 min CHX sample. Samples were then taken at 15, 30, 45, 60, and 90 min post-CHX for protein extraction and immunoblotting. To inhibit CK1δ or CK1ε; cells were treated with 1 µM PF670462 (Tocris Biosciences Cat# 3316) for 4 h prior to immunoblotting and immunofluorescence.

### Cell cycle arrest protocol

Cell cycle arrest protocol was adapted from Apraiz et al., 2017^45^. For both the S phase release and cell cycle arrest, U2OStx or U2OStx_CK1δ, were seeded onto four 10 cm dishes to 50% confluence. To release cells from S phase, a double thymidine (THY, final concentration of 2mM) was used as follows (see Fig. 5A): 24 h post-seeding, cells were treated with THY for 18 h, released into normal media for 6 h, treated with THY for a second block for 12 h. Cells were then released from the second THY block and sampled at 0, 1, and 2 h post-release.

To arrest cells in either G1/S or G2/M, either a double thymidine or a thymidine-nocodazole block, respectively, was used. For the unsynchronized sample, no additional treatment was performed (see Fig. 5C). For both the G1-and G2-arrested samples, after 24 h post-seeding, cells were treated with thymidine (THY; final concentration of 2 mM) for 18 h followed by a release into normal media for 6 h. To arrest cells in G1, cells were treated with a second thymidine block for 12 h. To arrest cells in G2, cells were treated with nocodazole (NOC, final concentration of 50 ng/mL) for 12 h. For the G2/M-arrested cells, an additional mitotic shake-off was performed to enrich the sample for cells in mid-mitosis that are loosely attached to the culture surface^44^. To induce CK1δ, DOX induction was performed on all samples 24 h prior to the end of either the double THY or THY-NOC treatments.

## Acknowledgements

We thank Alessia Ruggieri und Michael Knop for providing mKate2 and mNeonGreen templates, respectively. We also thank Cornelia Sala and Elmar Schiebel for the anti-PCNT antibody. This work was supported by the Deutsche Forschungsgemeinschaft, TRR186.

## Ethics declarations Competing interests

The authors declare no competing interests.

## Data Availability

The data supporting the findings of this study are available within the article. This study includes no data deposited in external repositories.

## Material Availability

Plasmids and cell lines generated in this study are available from the lead contact, Michael Brunner with a completed materials transfer agreement.

