## Supplementary material for "Cell cycle-coupled CK1δ turnover, autoinhibition, and activity": Figure 3 - Source data 1

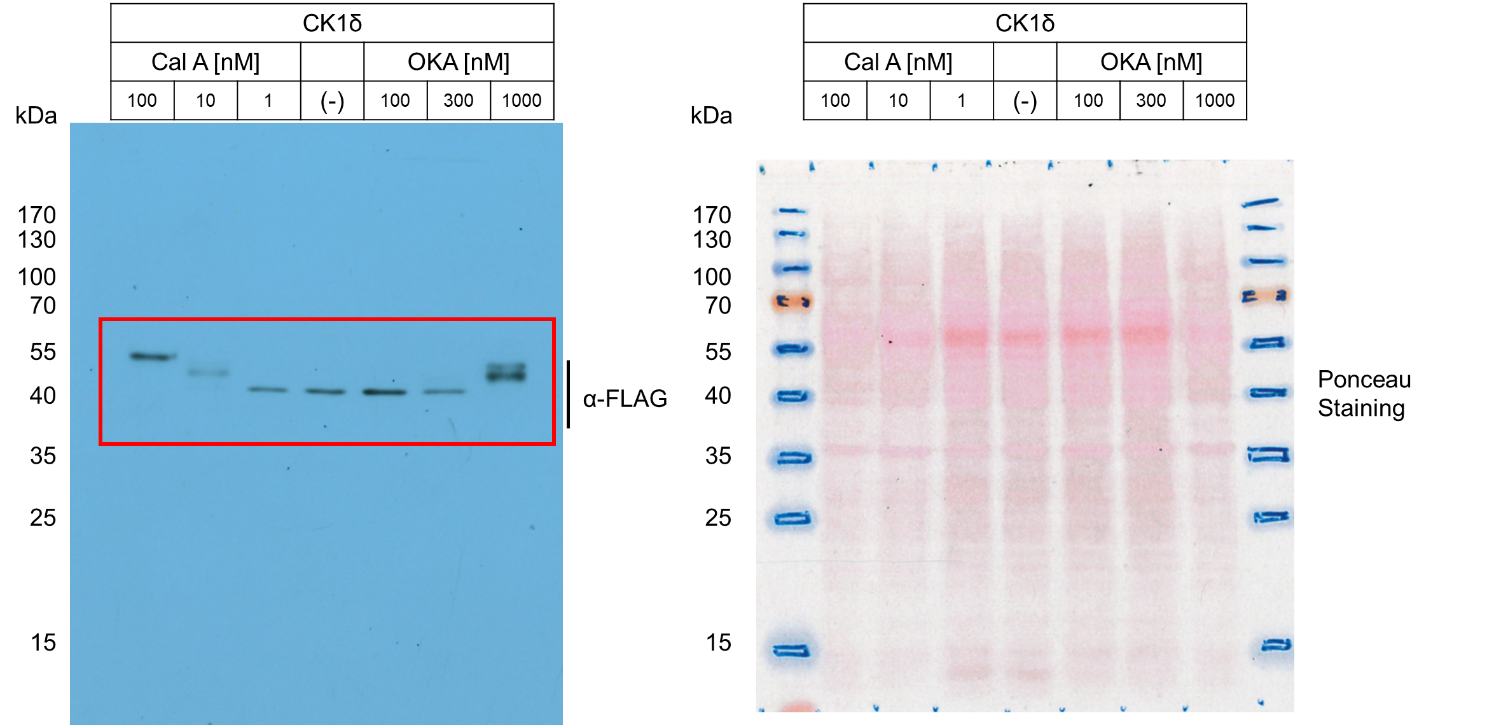

**Figure 3A – Source Data 1.** Left: Original film corresponding to Figure 3A. CK1δ-FLAG was induced and treated with either CalA or OKA at the corresponding dosages for 1h. Resulting blots were decorated with anti-FLAG antibody. Right: Original Ponceau Staining scan corresponding to Figure 3A. n = 3.

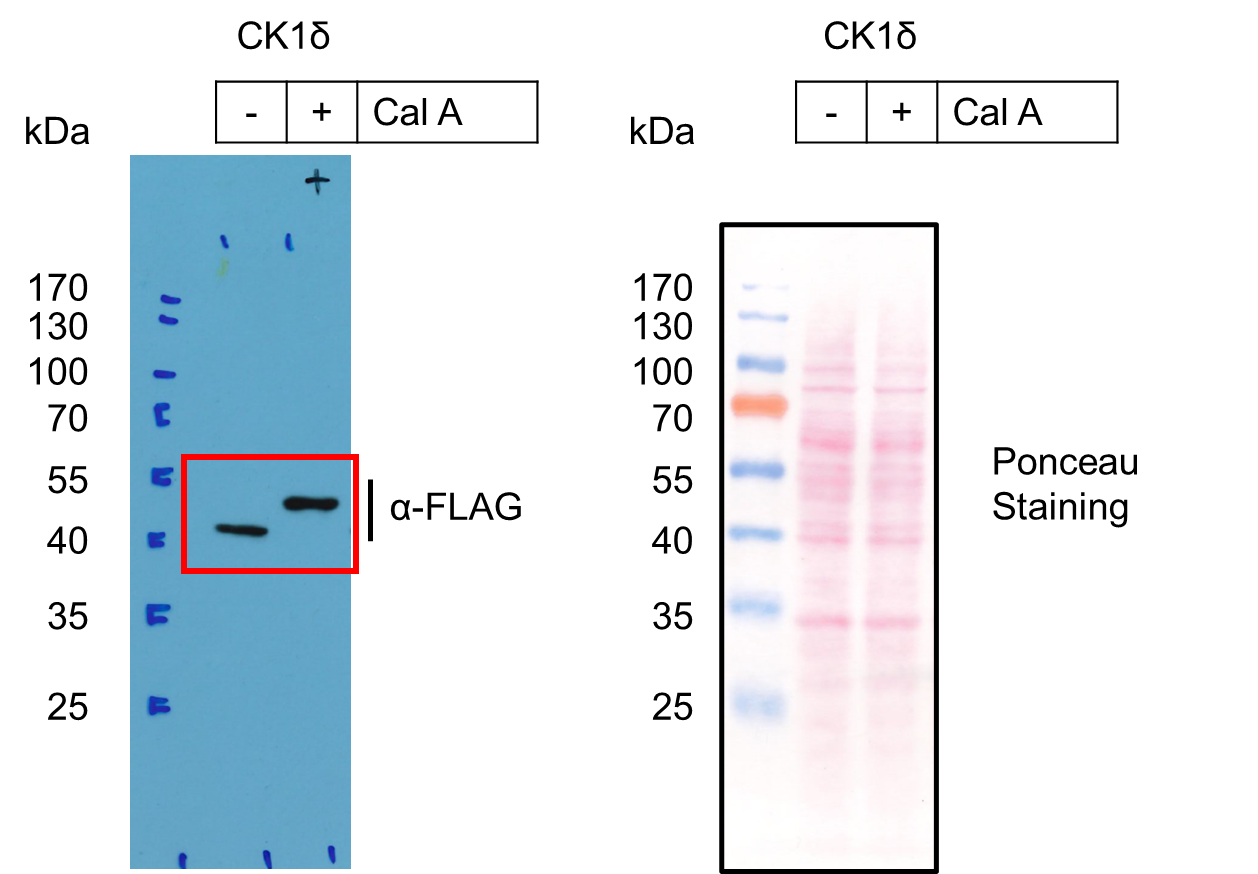

**Figure 3B – Source Data 1a.** Left: Original film corresponding to Figure 3B (left). CK1δ-FLAG was induced and treated with CalA. Resulting blots were decorated with anti-FLAG antibody. Right: Original Ponceau Staining scan corresponding to Figure 3B (left). n = 3.

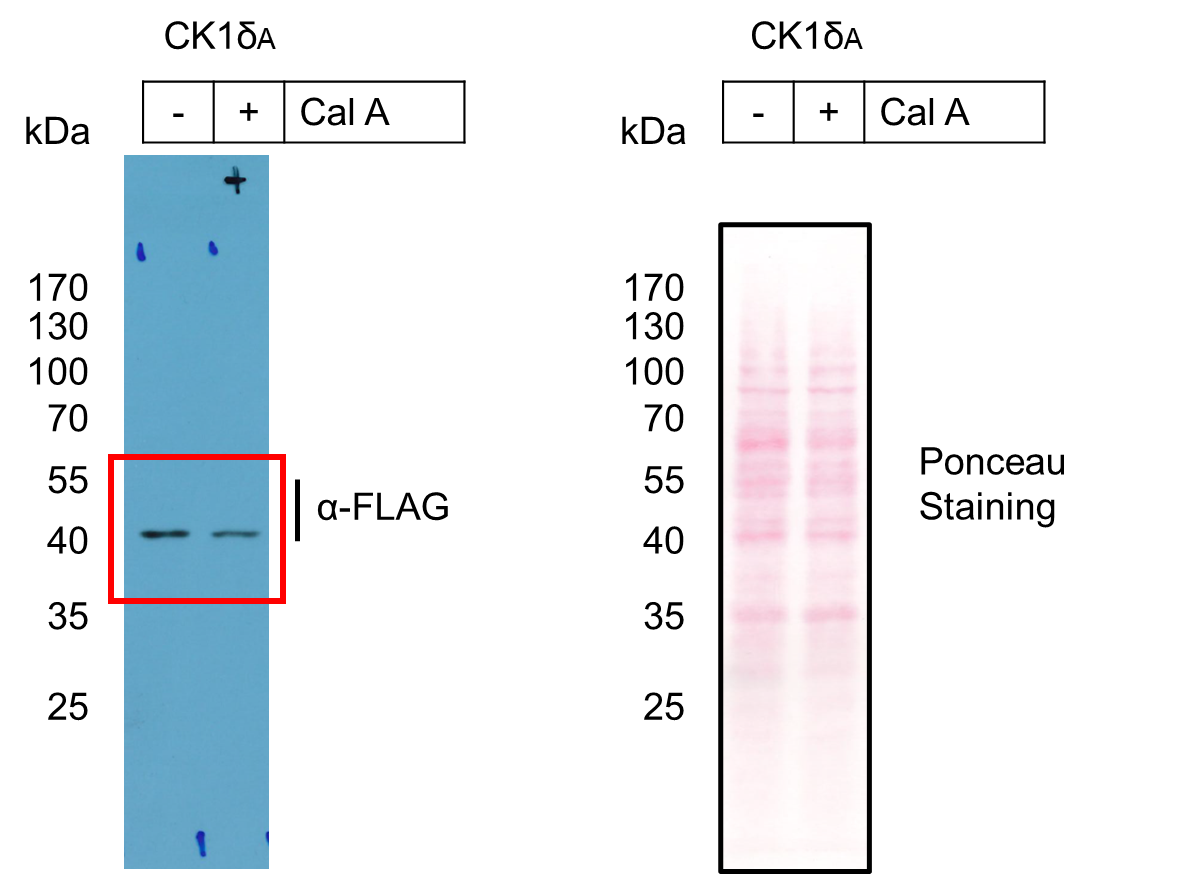

**Figure 3B – Source Data 1b.** Left: Original film corresponding to Figure 3B (right). CK1δ_A_-FLAG was induced and treated with CalA. Resulting blots were decorated with anti-FLAG antibody. Right: Original Ponceau Staining scan corresponding to Figure 3B (right). n = 3.

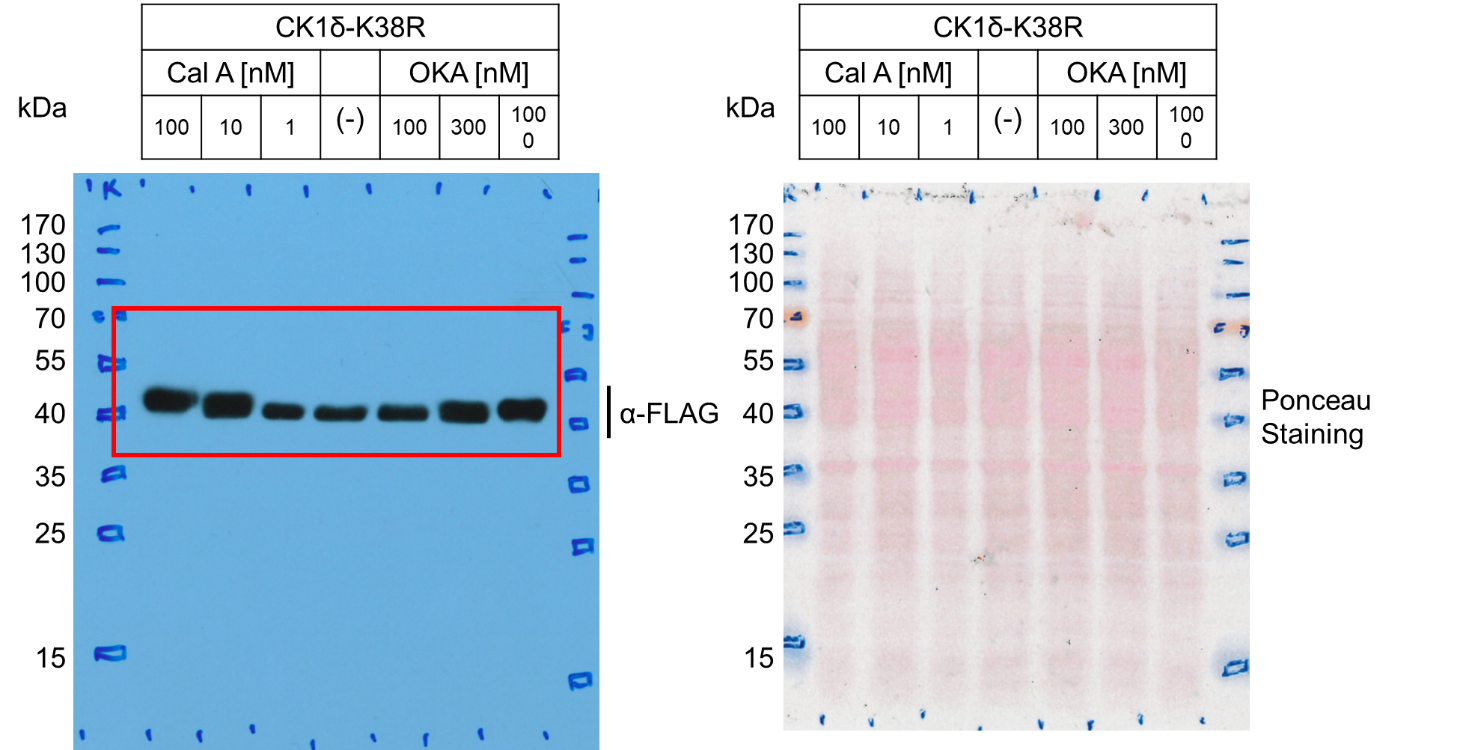

**Figure 3C – Source Data 1.** Left: Original film corresponding to Figure 3C. CK1δ-K38R-FLAG was induced and treated with either CalA or OKA at the corresponding dosages for 1h. Resulting blots were decorated with anti-FLAG antibody. Right: Original Ponceau Staining scan corresponding to Figure 3C. n = 3.

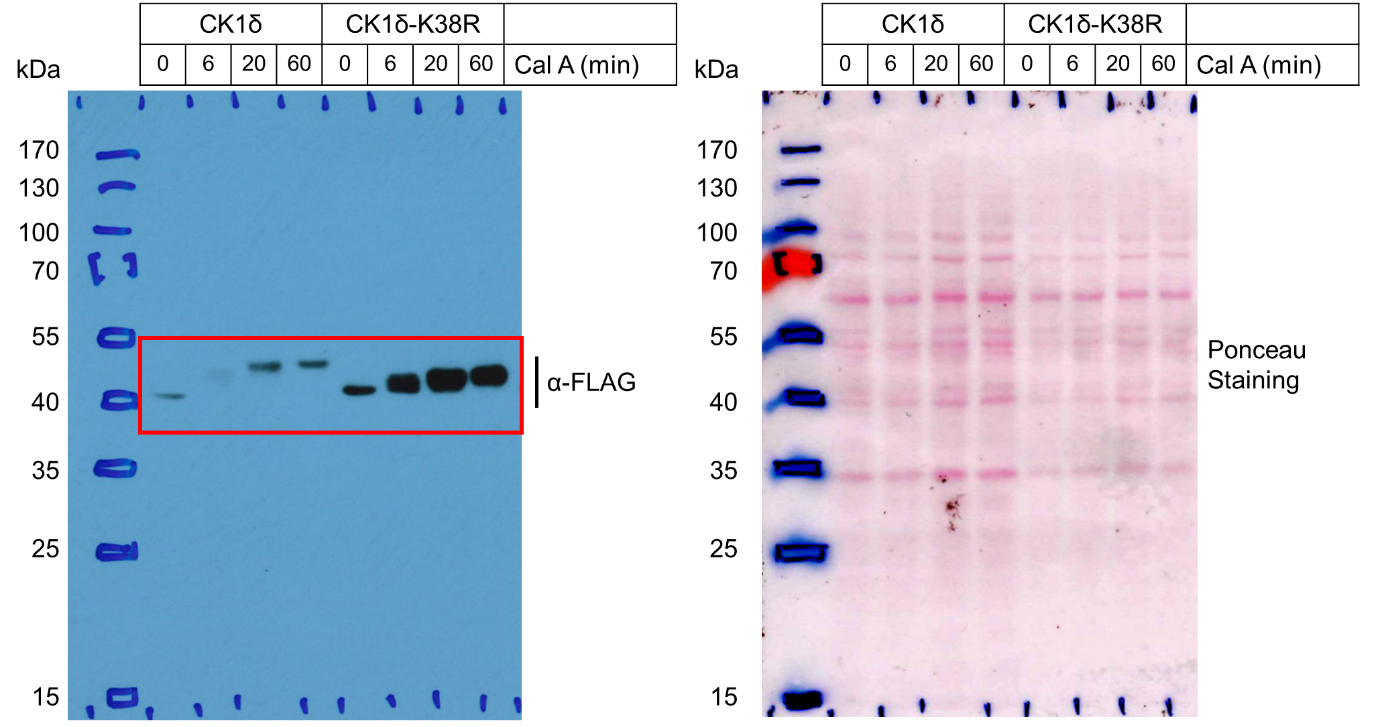

**Figure 3D – Source Data 1.** Left: Original film corresponding to Figure 3D. CK1δ and CK1δ-K38R were induced and treated with Cal A over the specified time-course. The resulting blot was decorated with anti-FLAG antibody. Right: Original Ponceau Staining scan corresponding to Figure 4D.  n = 3.
