## Supplementary figures and images for "Cell cycle-coupled CK1δ turnover, autoinhibition, and activity"

### Figure 3A_IB_CalA-OKA_CK1d_FLAG.tif

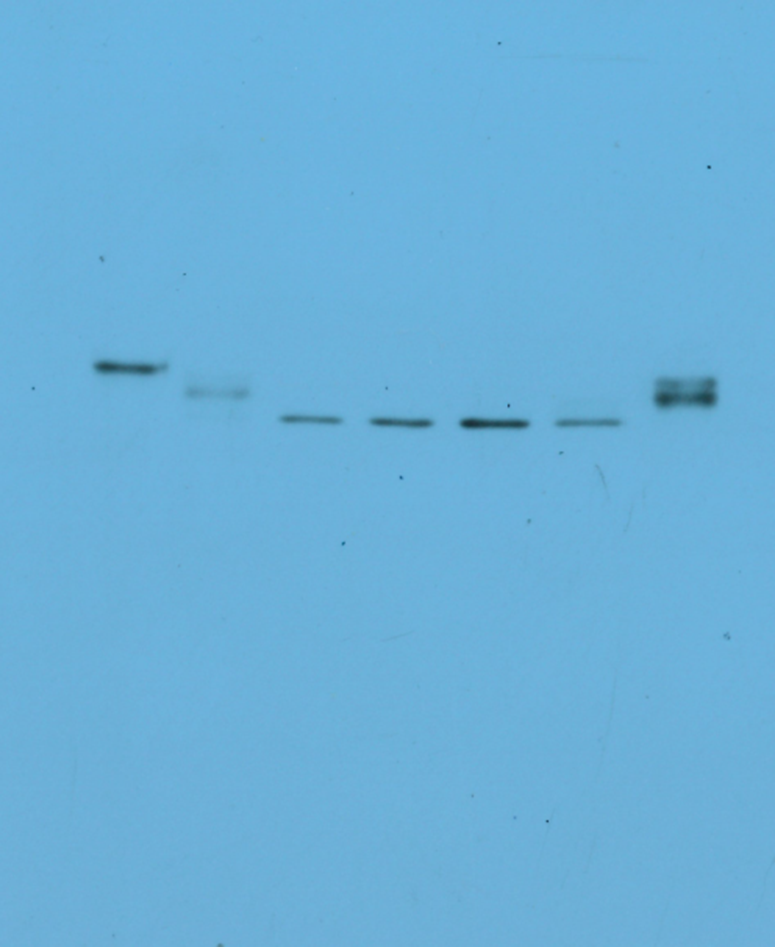

### Figure 3A_PonceauStain.tif

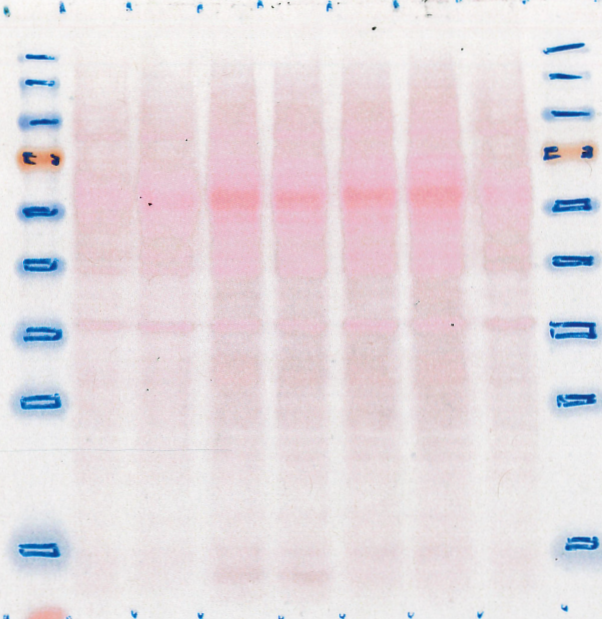

### Figure 3B_IB_CalA_CK1-CtermStoA_FLAG.tif

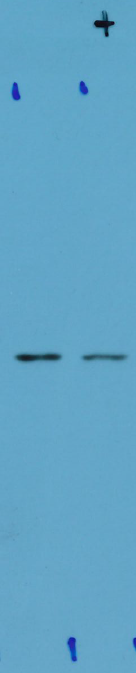

### Figure 3B_IB_CalA_CK1_FLAG.tif

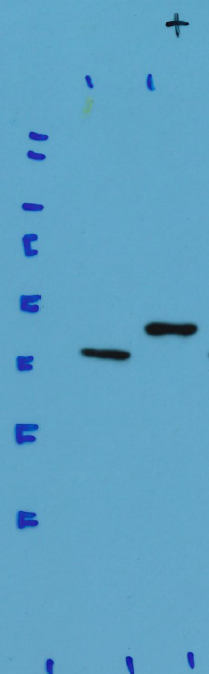

### Figure 3B_PonceauStain_CK1-CtermStoA.tif

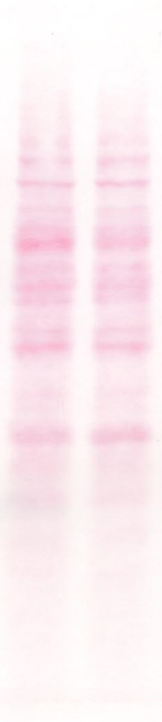

### Figure 3B_PonceauStain_CK1.tif

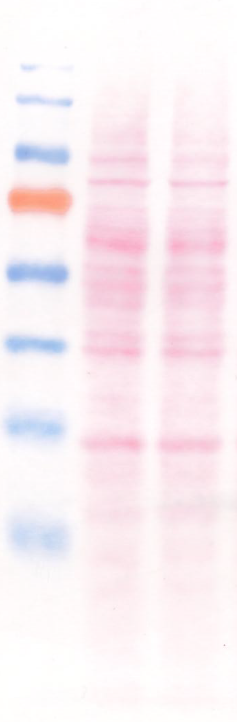

### Figure 3C_IB_CalA-OKA_CK1d-K38R_FLAG.tif

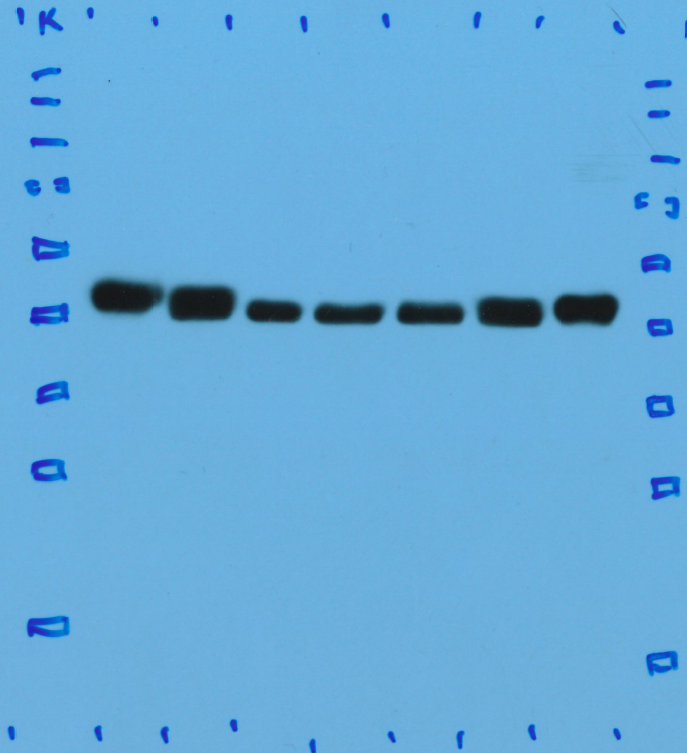

### Figure 3C_PonceauStain.tif

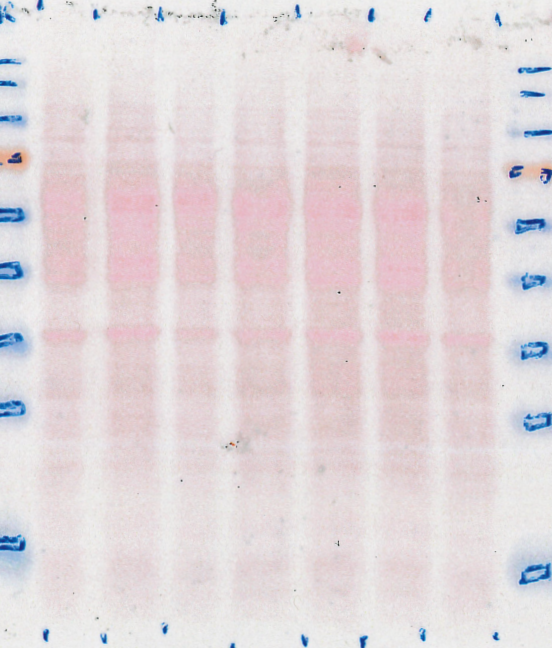

### Figure 3D_IB_CalATimeCourse_FLAG.tif

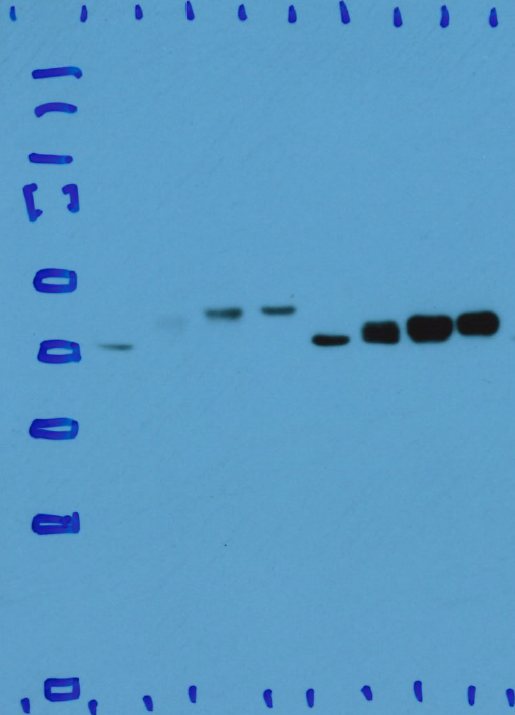

### Figure 3D_PonceauStain.tif

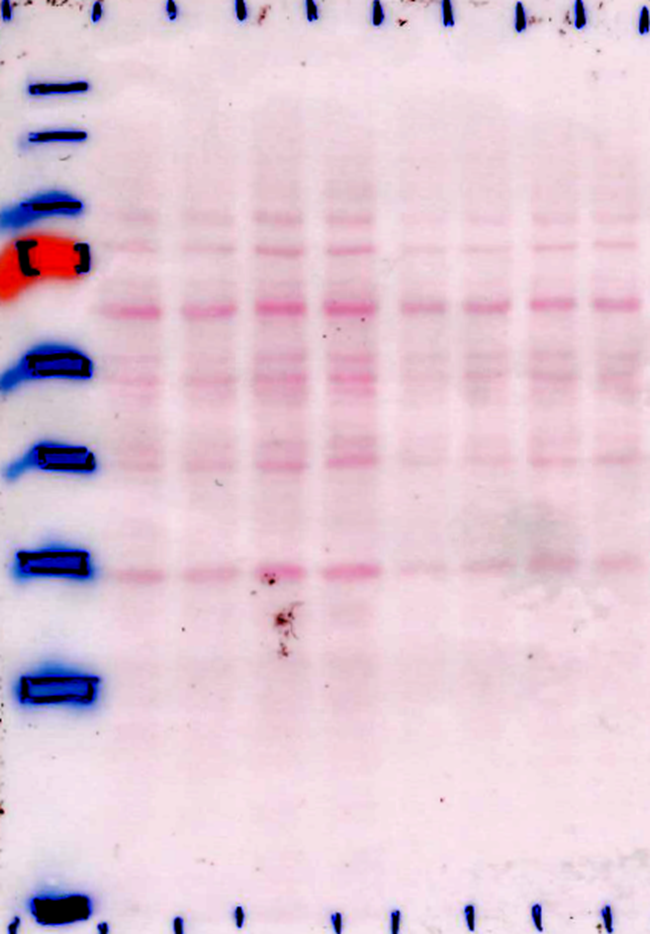

### Figure 4A_IB_CK1d_FLAG.tif

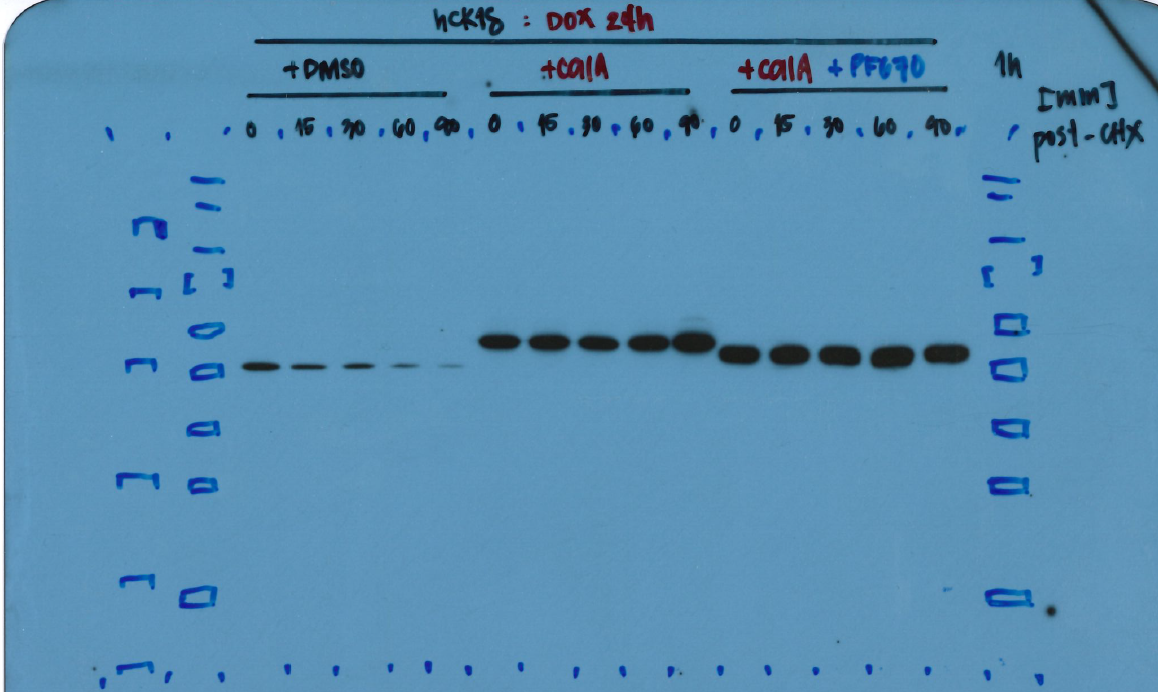

### Figure 4A_PonceauStaining.tif

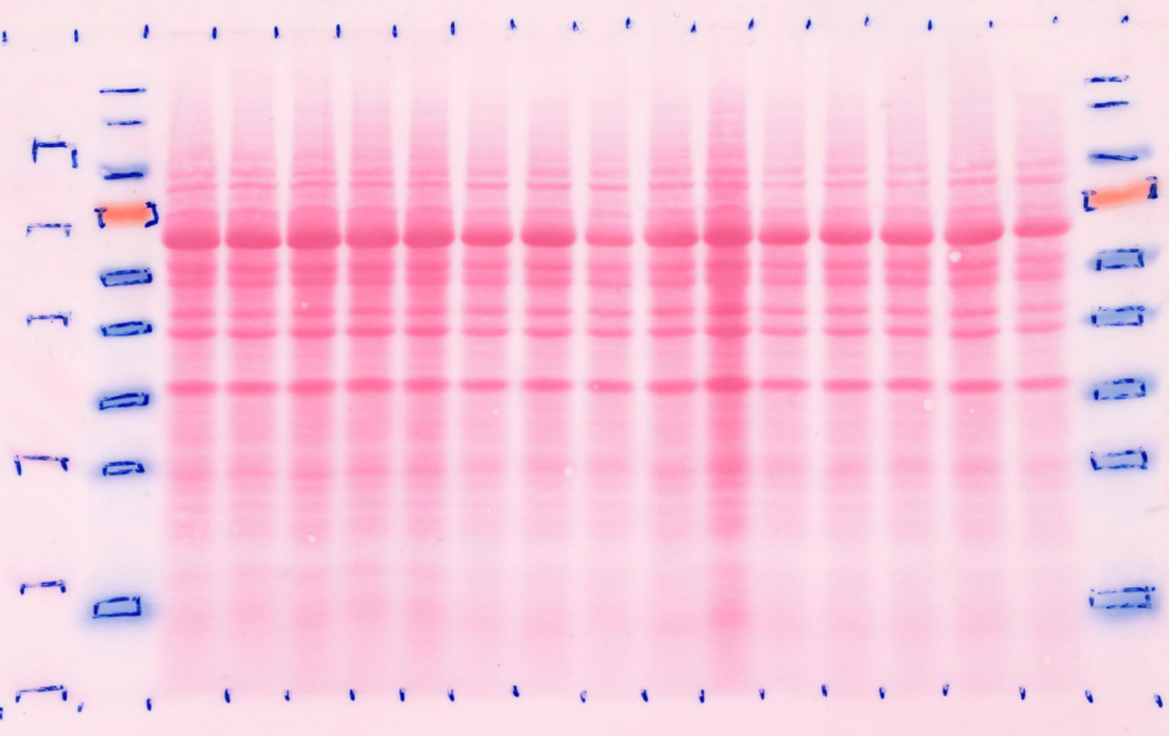

### Figure 4B_IB_CK1e_FLAG.tif

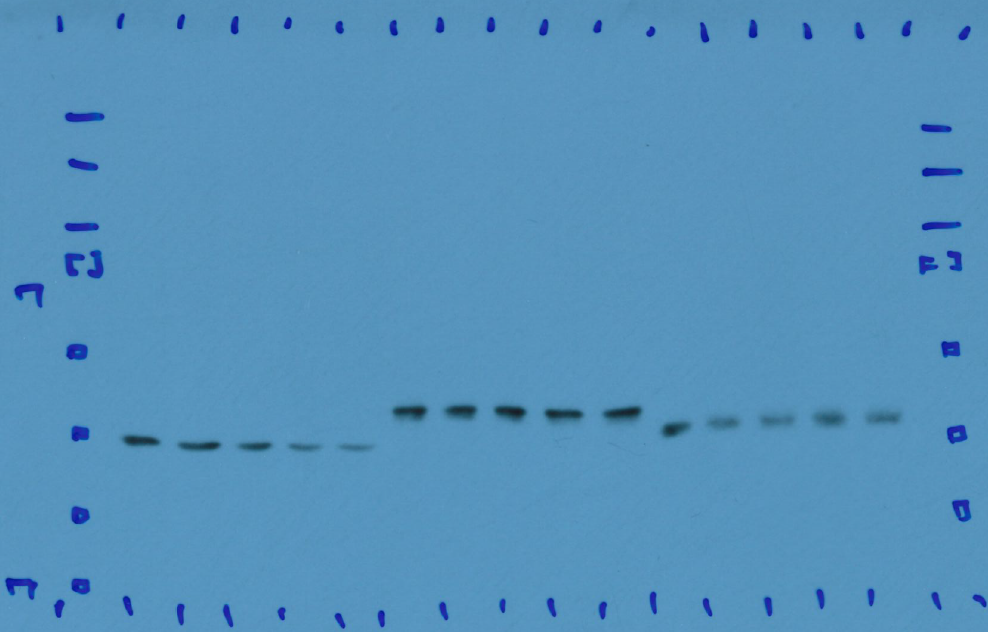

### Figure 4B_PonceauStaining.tif

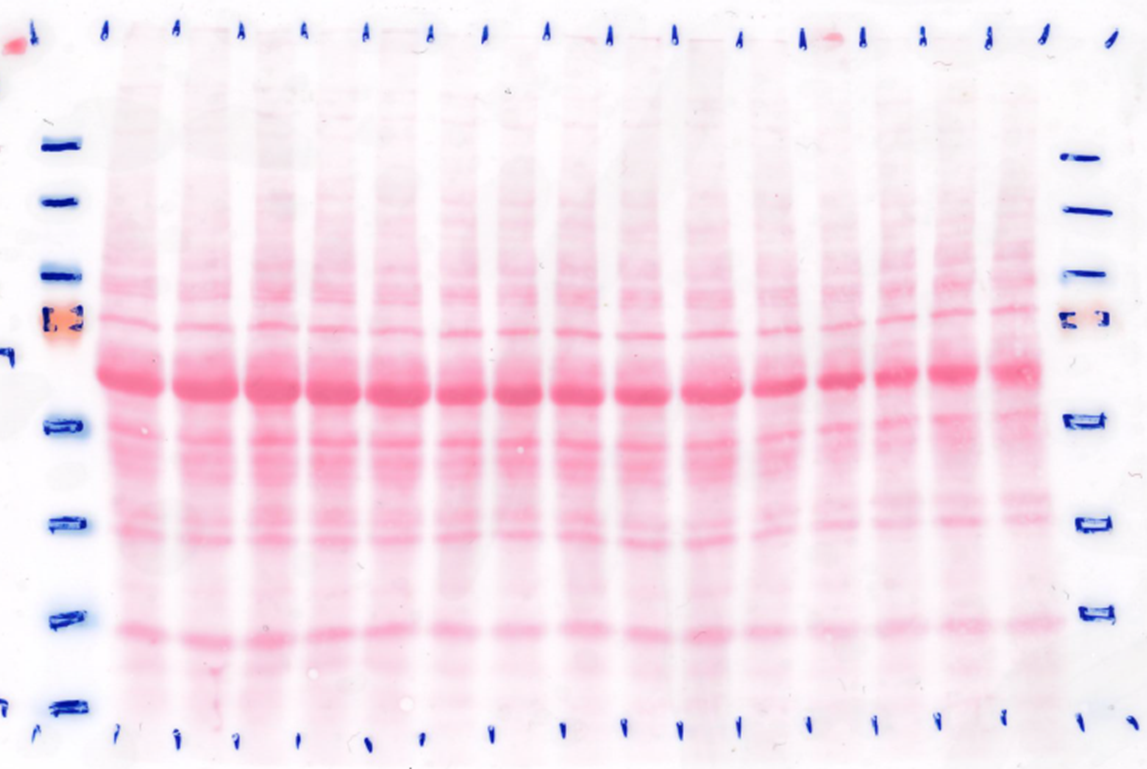

### Figure 4C_IB_NLS-CK1d_FLAG.tif

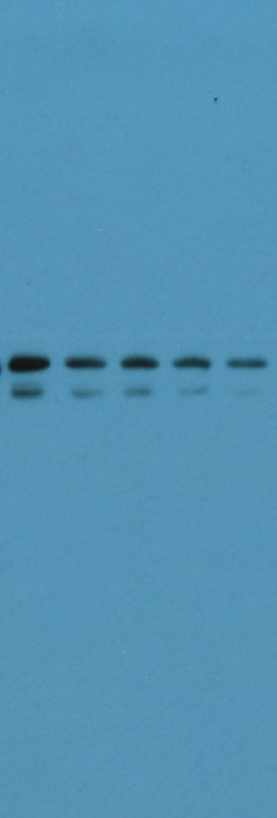

### Figure 4C_PonceauStaining.tif

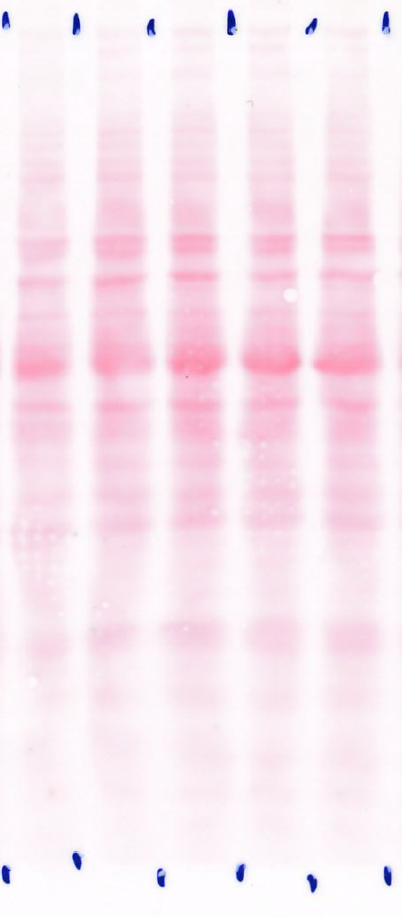

### Figure 4D_IB_NES-CK1d_FLAG.tif

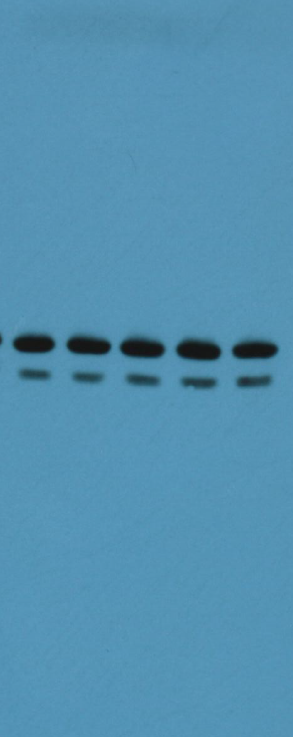

### Figure 4D_IB_PonceauStaining.tif

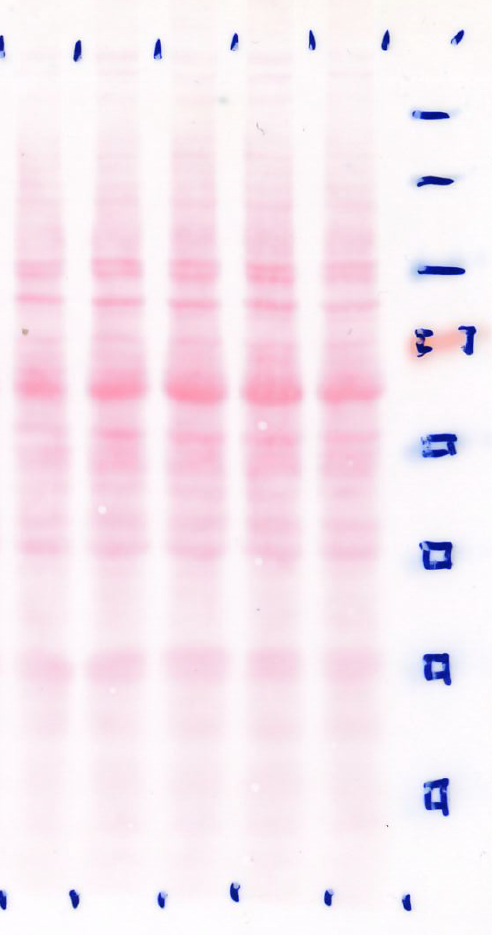

### Figure 5B_IB_post-S_FLAG.tif

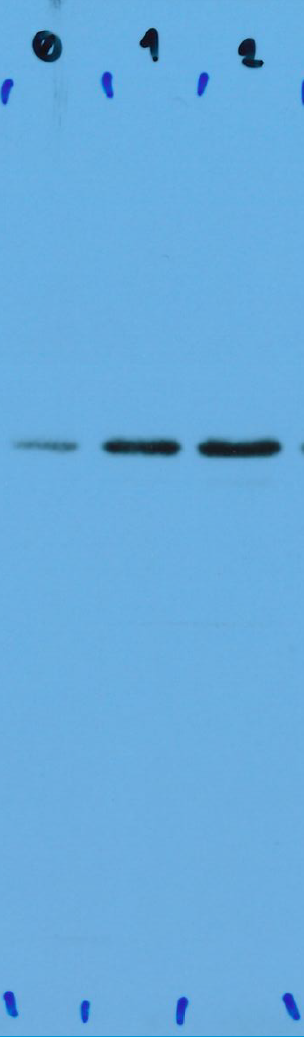

### Figure 5B_PonceauStaining.tif

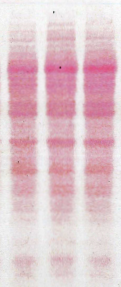

### Figure 5D_IB_CellCycleSync_oxCK1_FLAG.tif

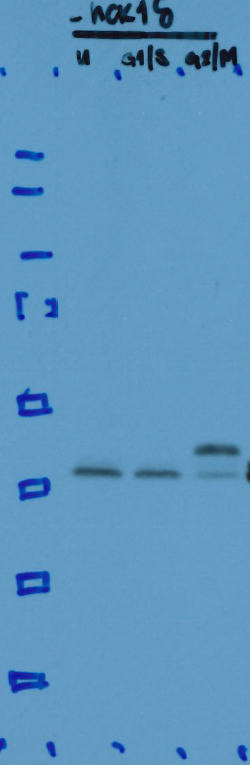

### Figure 5D_PonceauStaining.tif

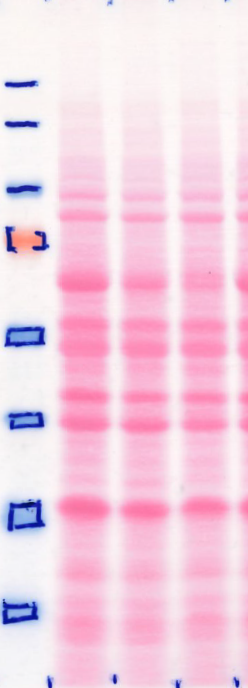

### Figure 5E_IB_CellCycleSync_CSNK1D.tif

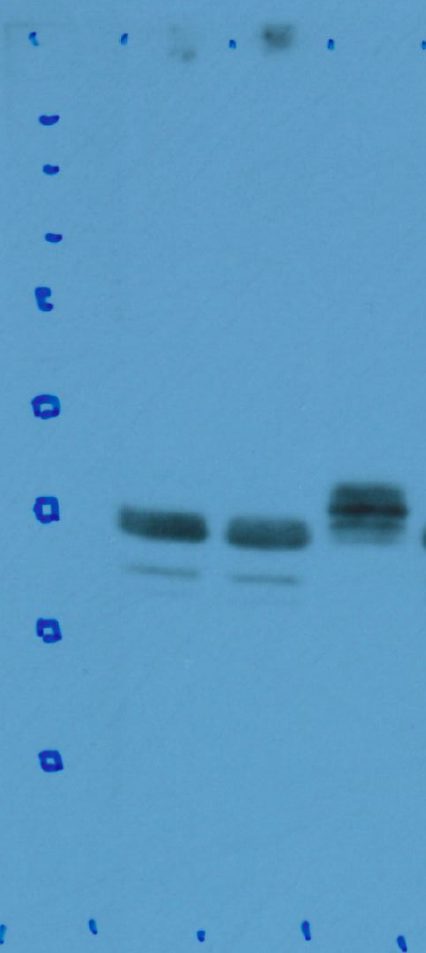

### Figure 5E_PonceauStaining.tif

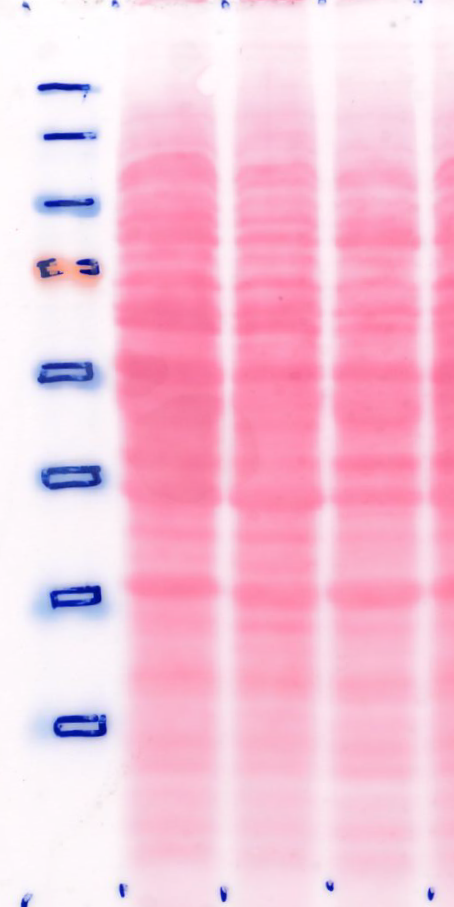
