## Supplementary material for "Cell cycle-coupled CK1δ turnover, autoinhibition, and activity": Figure 4 - Source data 1

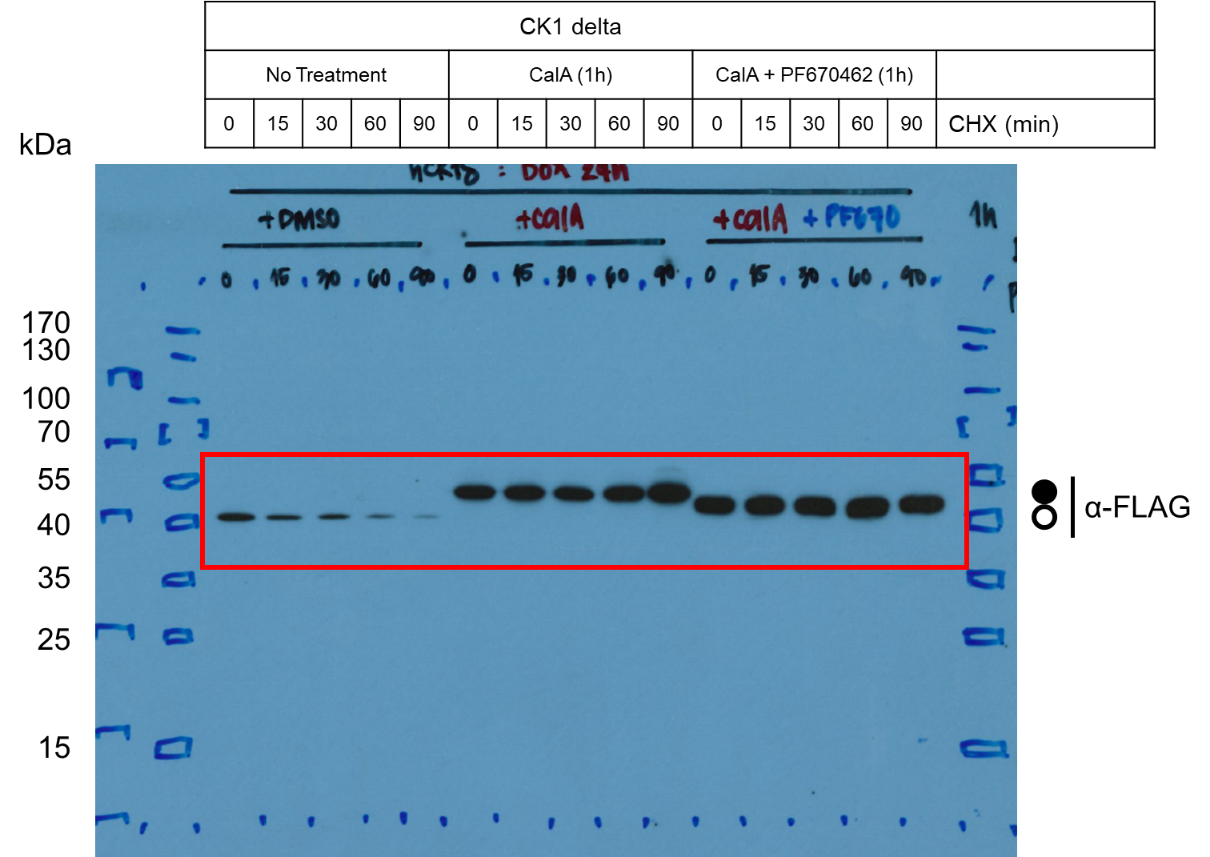

**Figure 4A – Source Data 1a.** Original film corresponding to Figure 4A. CK1δ-FLAG was induced and treated with a combination of CalA and PF670462 for 1h followed by CHX to arrest protein translation and assess the stability of the phosphorylated kinase. Circles correspond to hypo- (unfilled) and hyperphosphorylated (filled) kinase. Resulting blots were decorated with anti-FLAG antibody. n = 3.

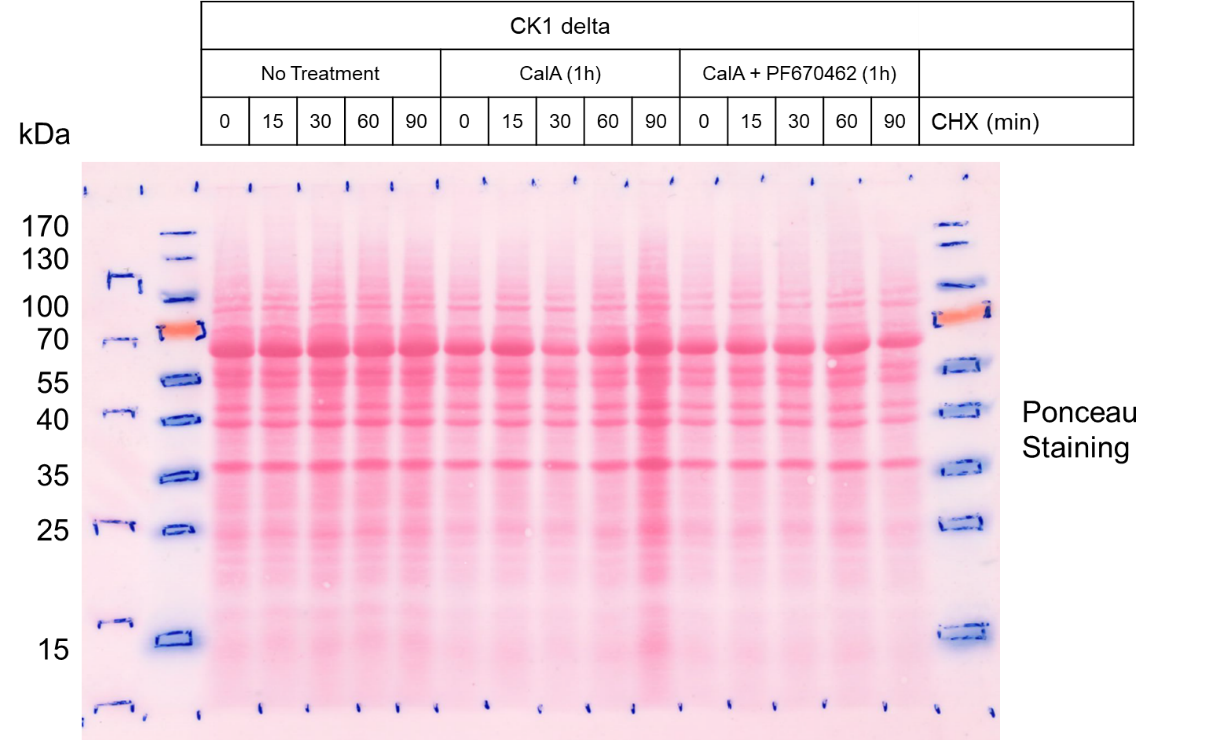

**Figure 4A – Source Data 1b.** Original Ponceau Staining scan corresponding to Figure 4A. n = 3.

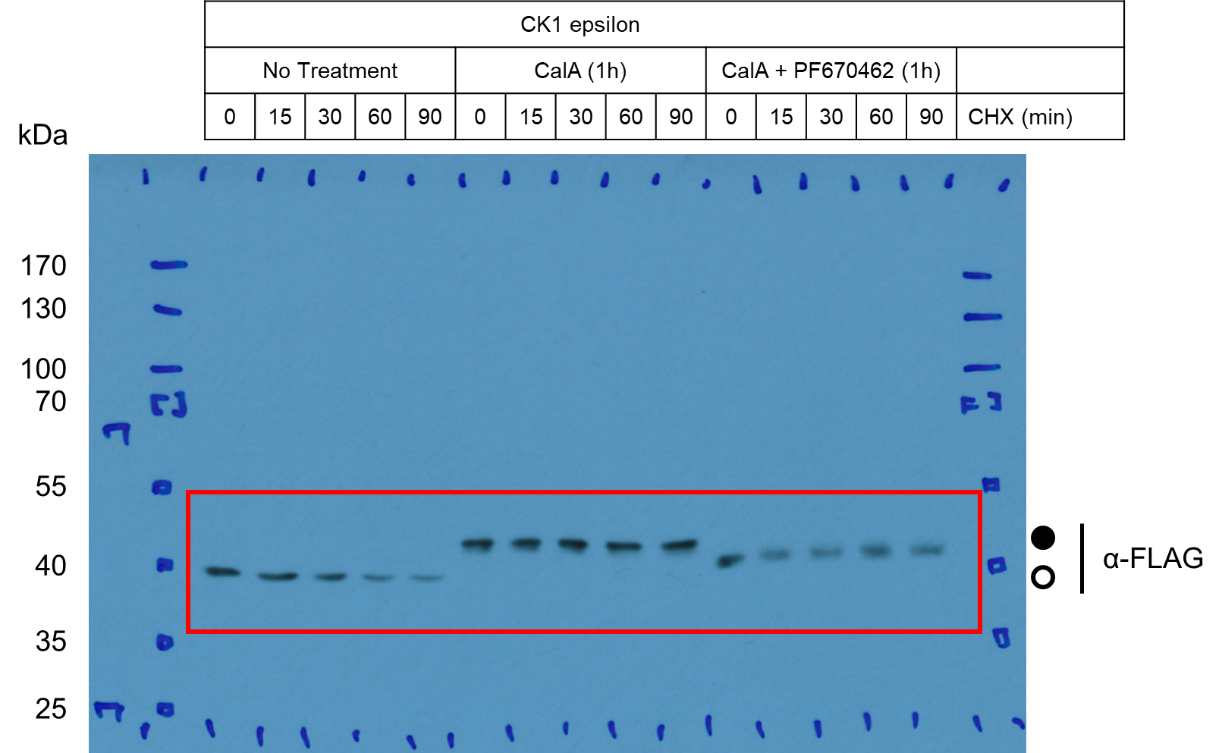

**Figure 4B – Source Data 1a.** Original film corresponding to Figure 4B. CK1ε-FLAG was induced and treated with a combination of CalA and PF670462 for 1h followed by CHX to arrest protein translation and assess the stability of the phosphorylated kinase. Circles correspond to hypo- (unfilled) and hyperphosphorylated (filled) kinase. Resulting blots were decorated with anti-FLAG antibody. n = 2.

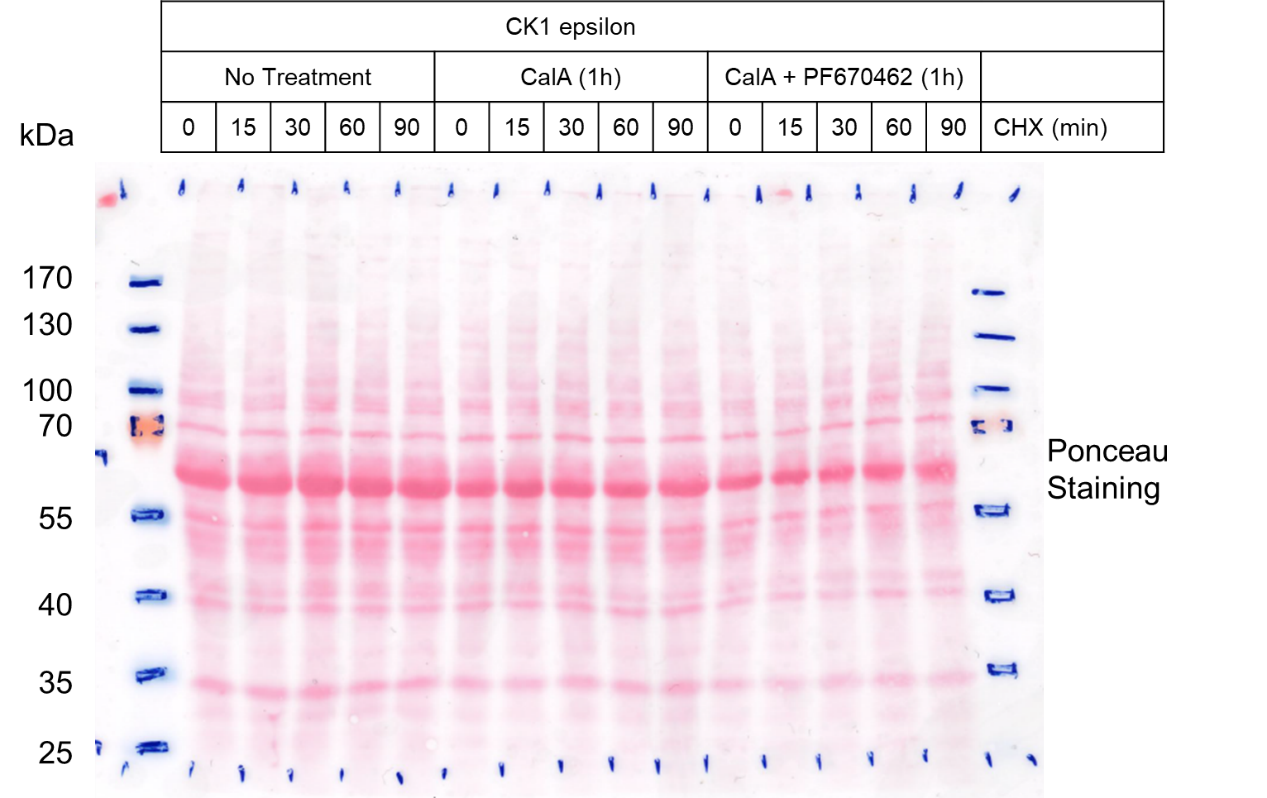

**Figure 4B – Source Data 1b.** Original Ponceau Staining scan corresponding to Figure 4B. n = 2.

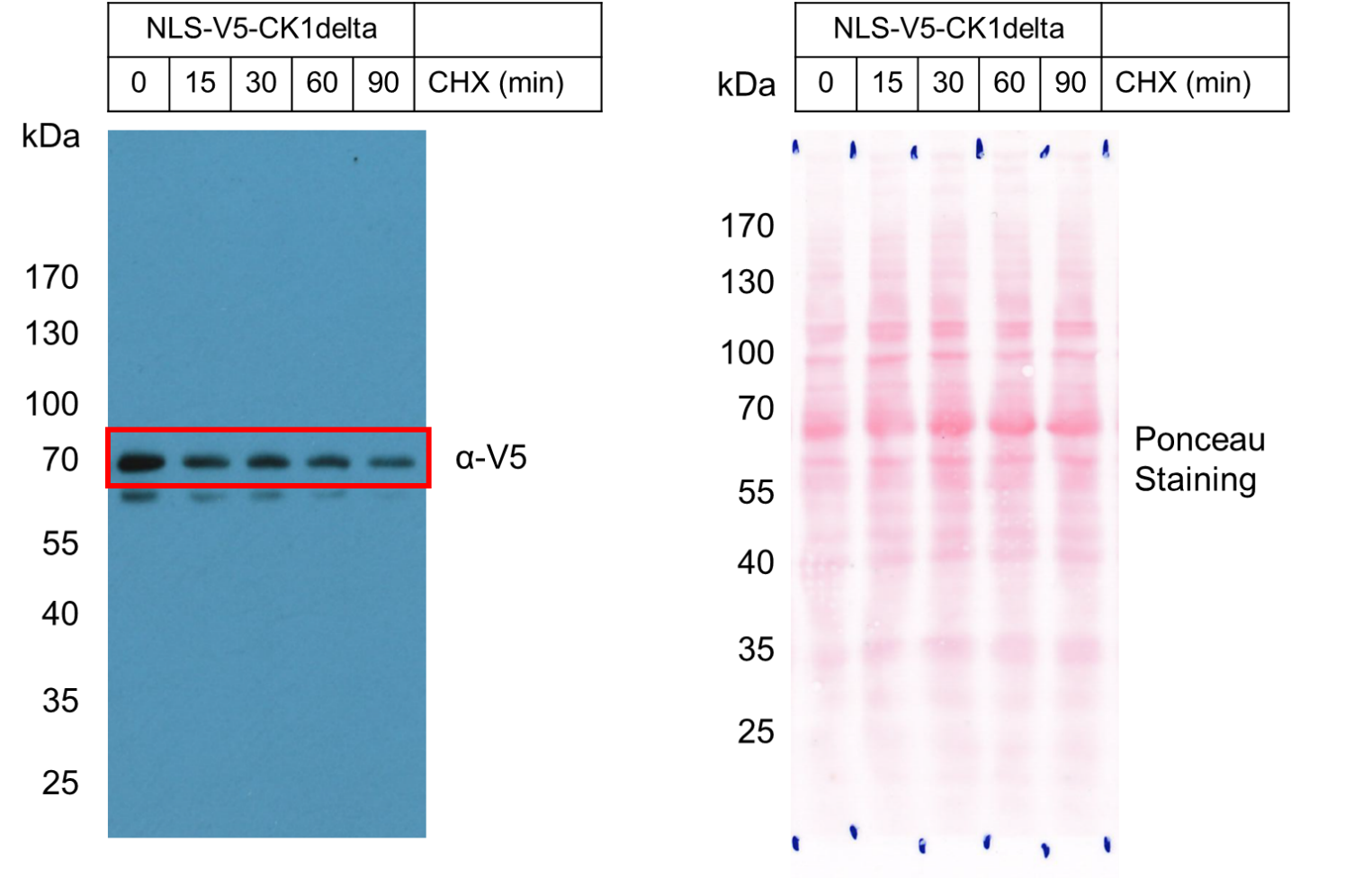

**Figure 4C – Source Data 1.** Left: Original film corresponding to Figure 4C. NLS-V5-CK1δ was induced and treated with CHX to arrest protein translation and assess the stability of the nuclear-localized kinase. Resulting blots were decorated with anti-V5 antibody. Right: Original Ponceau Staining scan corresponding to Figure 4C.  n = 3.

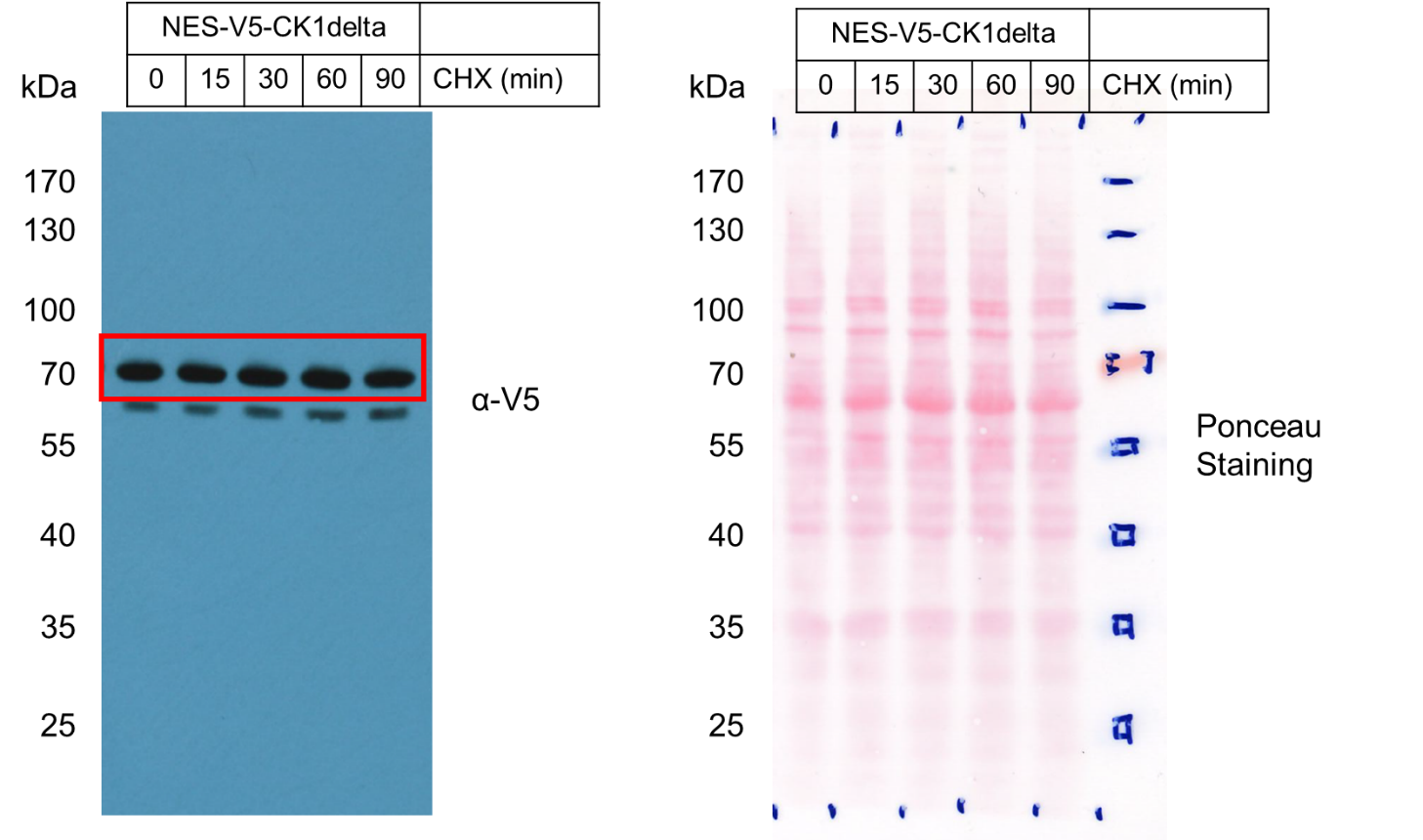

**Figure 4D – Source Data 1.** Left: Original film corresponding to Figure 4D. NES-V5-CK1δ was induced and treated with CHX to arrest protein translation and assess the stability of the cytoplasm-localized kinase. Resulting blots were decorated with anti-V5 antibody. Right: Original Ponceau Staining scan corresponding to Figure 4D.  n = 3.
