## Supplementary material for "Cell cycle-coupled CK1δ turnover, autoinhibition, and activity": Figure 5 - Source data 1

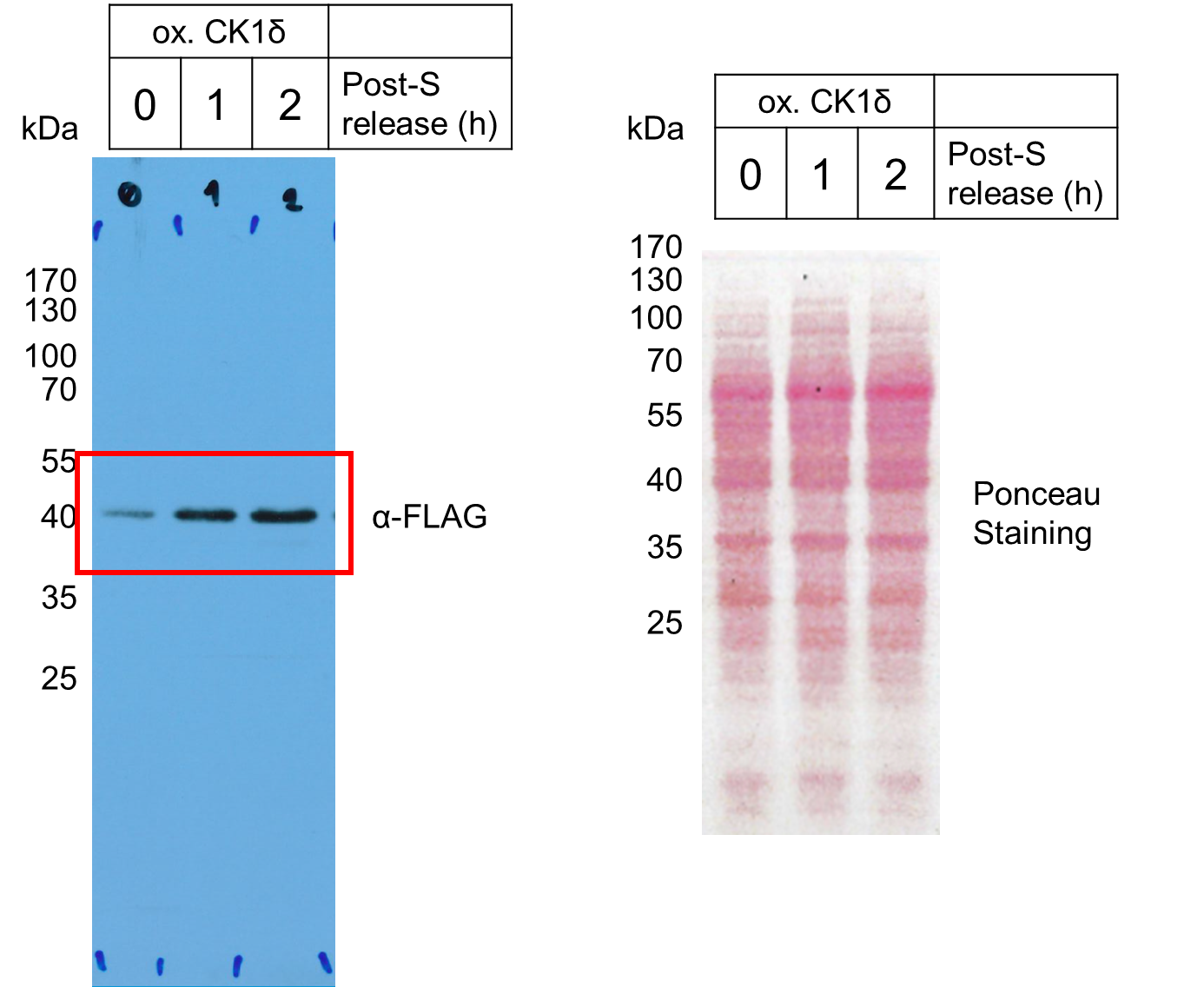

**Figure 5B – Source Data 1.** Left: Original film corresponding to Figure 5B. CK1δ-FLAG was induced followed by a double thymidine block and release and sampled as shown on Figure 5A. Resulting blots were decorated with anti-FLAG antibody. Right: Original Ponceau Staining scan corresponding to Figure 5B. n = 3.

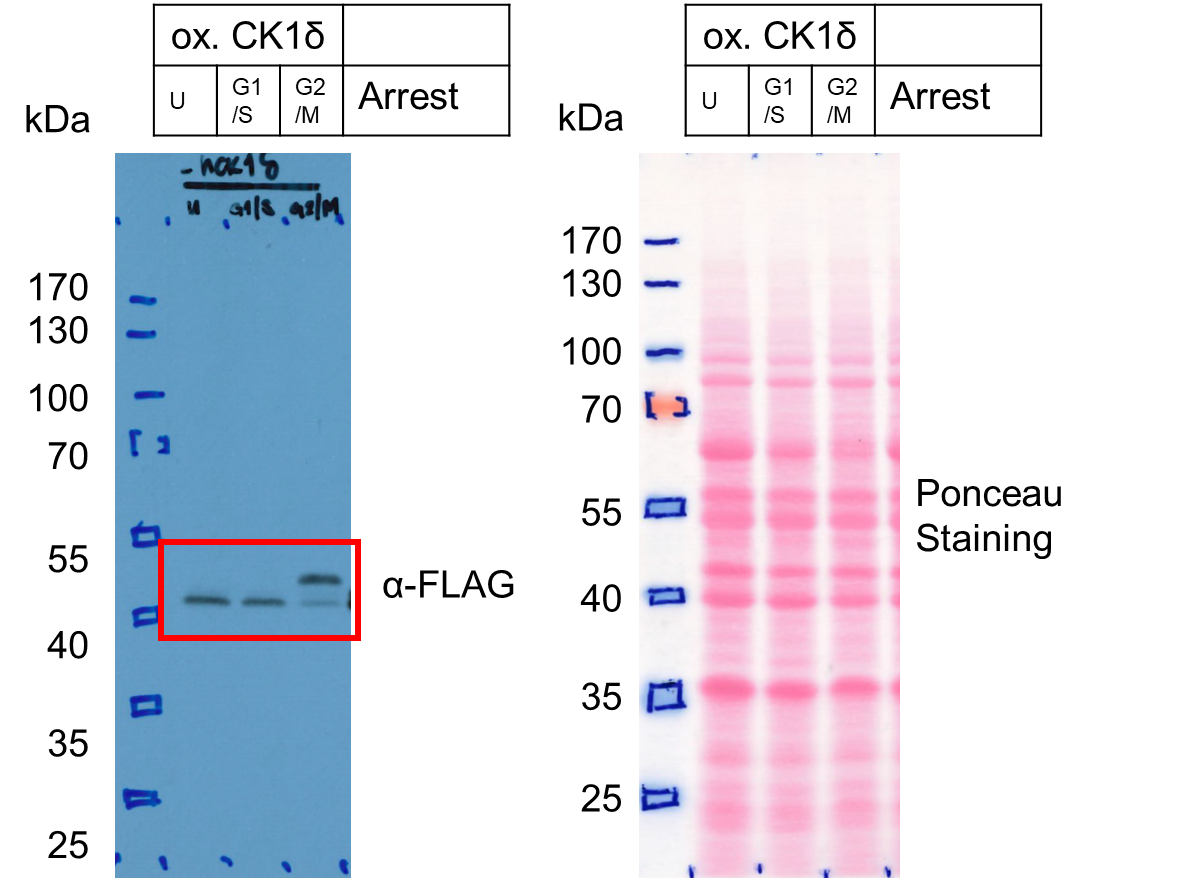

**Figure 5D – Source Data 1.** Left: Original film corresponding to Figure 5D. CK1δ-FLAG was induced followed by a cell cycle arrest protocol and sampled as shown on Figure 5C. Resulting blots were decorated with anti-FLAG antibody. Right: Original Ponceau Staining scan corresponding to Figure 5D. n = 3.

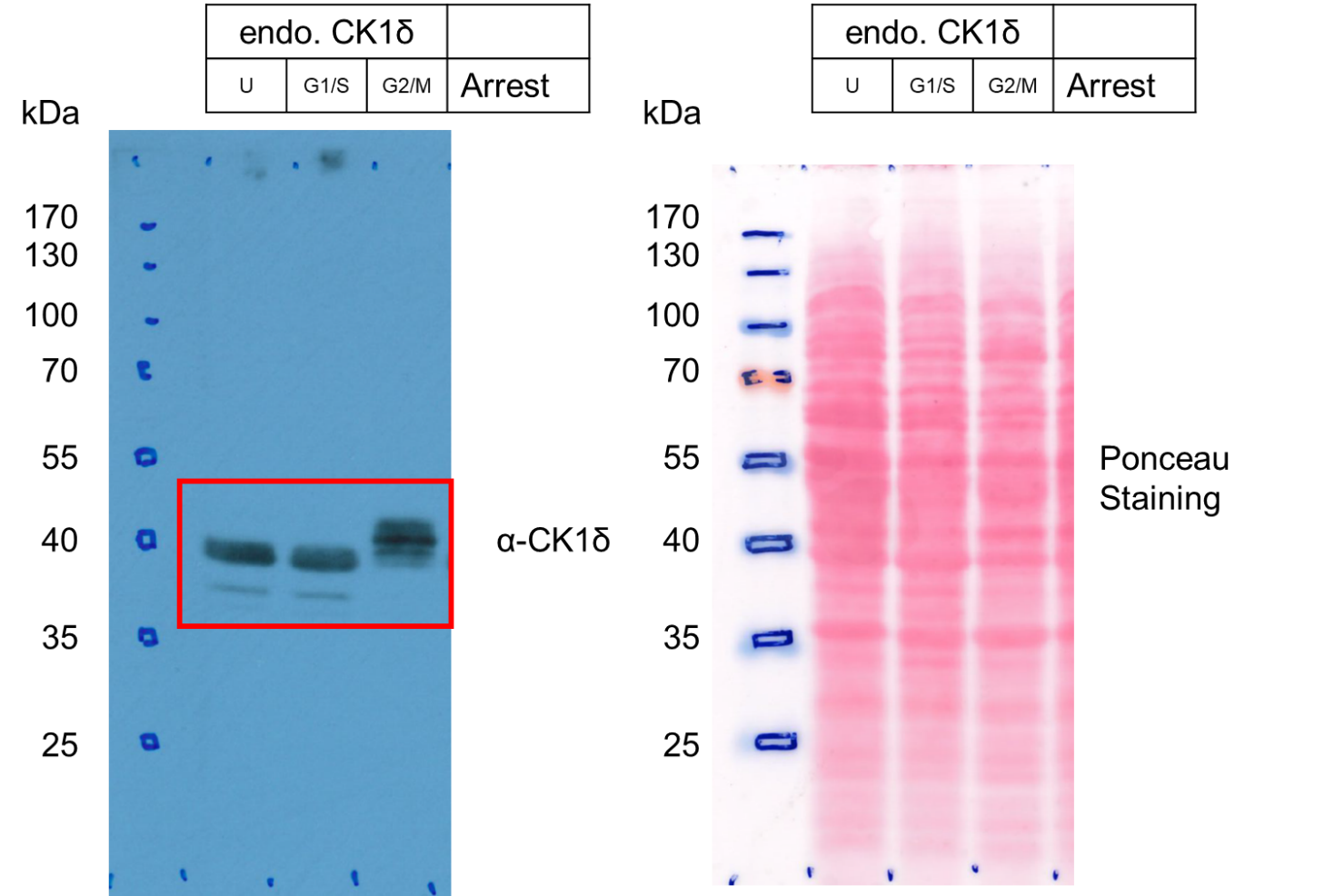

**Figure 5E – Source Data 1.** Left: Original film corresponding to Figure 5E. U2OStx cells were treated with a cell cycle arrest protocol and sampled as shown on Figure 5C. Resulting blots were decorated with anti-CK1δ antibody. Right: Original Ponceau Staining scan corresponding to Figure 5E. n = 3.
